# ACE2 engagement exposes the fusion peptide to pan-coronavirus neutralizing antibodies

**DOI:** 10.1101/2022.03.30.486377

**Authors:** Jun Siong Low, Josipa Jerak, M. Alejandra Tortorici, Matthew McCallum, Dora Pinto, Antonino Cassotta, Mathilde Foglierini, Federico Mele, Rana Abdelnabi, Birgit Weynand, Julia Noack, Martin Montiel-Ruiz, Siro Bianchi, Fabio Benigni, Nicole Sprugasci, Anshu Joshi, John E. Bowen, Alexandra C. Walls, David Jarrossay, Diego Morone, Philipp Paparoditis, Christian Garzoni, Paolo Ferrari, Alessandro Ceschi, Johan Neyts, Lisa A. Purcell, Gyorgy Snell, Davide Corti, Antonio Lanzavecchia, David Veesler, Federica Sallusto

## Abstract

Coronaviruses use diverse Spike (S) glycoproteins to attach to host receptors and fuse with target cells. Using a broad screening approach, we isolated from SARS-CoV-2 immune donors seven monoclonal antibodies (mAbs) that bind to all human alpha and beta coronavirus S proteins. These mAbs recognize the fusion peptide and acquire high affinity and breadth through somatic mutations. Despite targeting a conserved motif, only some mAbs show broad neutralizing activity in vitro against alpha and beta coronaviruses, including Omicron BA.1 variant and bat WIV-1, and reduce viral titers and pathology in vivo. Structural and functional analyses show that the fusion peptide-specific mAbs bind with different modalities to a cryptic epitope which is concealed by prefusion-stabilizing ‘2P’ mutations and becomes exposed upon binding of ACE2 or ACE2-mimicking mAbs. This study identifies a new class of pan-coronavirus neutralizing mAbs and reveals a receptor-induced conformational change in the S protein that exposes the fusion peptide region.

---

Human-infecting coronaviruses (HCoVs) are distributed across two genera: *alphacoronavirus*, which includes NL63 and 229E, and *betacoronavirus*, which includes OC43, HKU1, SARS-CoV-2, SARS-CoV and MERS-CoV. The spike (S) glycoprotein, which facilitates viral entry into host cells via ACE2 receptor, is composed of the S_1_ and S_2_ subunits, is highly divergent, with only ∼30% sequence identity between alpha and beta coronaviruses and is the main target of neutralizing antibodies (*1–3*). Previous studies have described neutralizing monoclonal antibodies (mAbs) that cross-react amongst sarbecoviruses by targeting the receptor binding domain (RBD) (*4–10*) or more broadly across beta coronaviruses by targeting the stem helix (*11–15*). However, neutralizing mAbs that target both the alpha and beta coronaviruses have not been reported. Broadly neutralizing mAbs, as exemplified for those targeting influenza viruses or HIV-1 (*16–21*), can potentially be used for prophylaxis or therapy and to guide the design of vaccines eliciting broadly protective immunity (*22, 23*).

## Isolation of pan-coronavirus mAbs from convalescent and vaccinated individuals

To search for rare antibodies that cross-react with alpha and beta coronaviruses, we stimulated under limiting conditions, total peripheral blood mononuclear cells (PBMCs) from SARS-CoV-2 immune donors, in the presence of R848 and IL-2, which selectively induce the proliferation and differentiation of memory B cells (*24*). On day 12, the specificities of IgGs secreted in the culture supernatants were tested by ELISA against a panel of recombinant S proteins from beta and alpha HCoVs (**Fig. 1, A** to **C**). The number of SARS-CoV-2 IgG-positive cultures was generally higher in COVID-19 convalescent patients and in SARS-CoV-2 vaccinees with prior infection (pre-immune) as compared to vaccinees without prior infection (naïve). Most SARS-CoV-2 IgG-positive cultures were either monospecific or cross-reactive with the closely related SARS-CoV, while a small fraction was also reactive with OC43 and HKU1 (**Fig. 1, A** to **D**), consistent with previous serological analyses (*25–27*). Remarkably, six cultures (out of >4,000) from 5 individuals (out of 43) cross-reacted with all alpha and beta HCoV S proteins tested (**Fig. 1, A** to **D**), suggesting that memory B cells producing pan-coronavirus antibodies might exist at very low frequency.

**Fig. 1.**
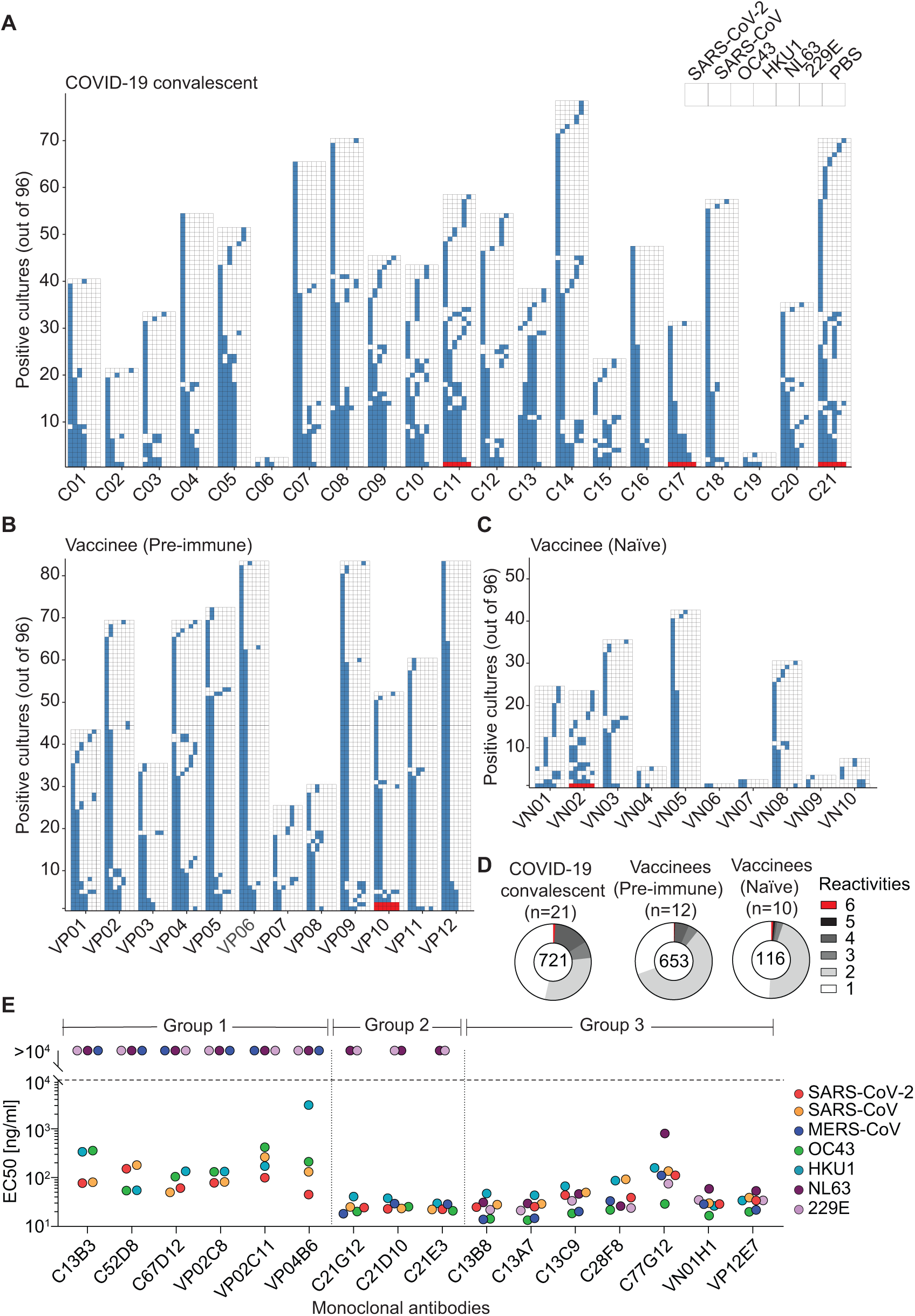
Rare pan-reactive memory B cells are elicited upon natural infection or vaccination. (**A** to **D**) Total PBMCs from COVID-19 convalescents (C) (**A**), vaccinees with prior infection (pre-immune) (VP) (**B**), and vaccinees without prior infection (naïve) (VN) (**C**) were plated in replicate 96 wells (10^4^ cells/well) and stimulated with TLR agonist R848 (2.5 µg/ml) in the presence of IL-2 (1,000 U/ml). Twelve days later, the supernatant of each culture was screened, in parallel, for the specificities of the secreted IgG antibodies to recombinant S proteins from SARS-CoV-2, SARS-CoV, OC43, HKU1, NL63 and 229E by ELISA. The skyline plot provides a detailed view of the specificities of each culture well (represented in rows) to the respective antigens (in subcolumn) from each donor (in column). The order of the antigens is indicated in the legend and uncoated plates (PBS) were used as control. If OD 405nm value exceeds the cut-off value determined by average OD 405nm of PBS wells + 4*standard deviation, the culture was considered reactive to the antigen and indicated with colored cells. Only cultures exhibiting reactivity to at least one antigen are shown. Red cells highlight the six cultures exhibiting reactivity to all alpha and beta HCoV S proteins tested. (**D**) Pie chart shows the cross-reactivity patterns of SARS-CoV-2 S positive cultures from **A** to **C**. The total number of SARS-CoV-2 S-positive cultures from each cohort is indicated at the centre of the pie. (**E**) Using a two-step screening strategy as depicted in Fig. S1, 16 monoclonal antibodies (mAbs) that cross-reacted with multiple HCoV S proteins were isolated from 11 donors (C series from convalescent donors, VP and VN series from pre-immune and naïve vaccinees). Shown are EC50 values to the respective S protein antigens, measured by ELISA. mAbs are grouped based on the reactivity patterns. Shown is one representative experiment out of at least 2 performed. Group 2 mAbs C21G12, C21D10, and C21E3 were described in a separate study (*11*) (designated P34G12, P34D10, and P34E3, respectively).

To isolate pan-coronavirus mAbs, we combined the broad screening of polyclonally-activated memory B cells with the sorting and cloning of antibody secreting cells to retrieve paired heavy and light chain sequences (**Fig. S1**). Using this approach, we isolated 16 SARS-CoV-2-S-specific mAbs that cross-reacted with various HCoV S proteins **(Fig. 1E** and **Fig. S2A**). Six mAbs (Group 1) cross-reacted with SARS-CoV, OC43 and HKU1 S proteins and bound with EC50 values ranging from 45 ng/ml to 3,000 ng/ml. Three mAbs (Group 2) cross-reacted with all beta HCoV S proteins with high-avidity, as illustrated by their EC50 values ranging from 18 ng/ml to 40 ng/ml (**Fig. 1E**). These mAbs were found to target the stem helix region and were described in a separate study (*11*). Remarkably, the remaining seven mAbs (Group 3) exhibited the broadest cross-reactivity to both alpha and beta HCoVs with EC50 values ranging from 29 ng/ml to 800 ng/ml (**Fig. 1E**). These pan-reactive mAbs, which will be the focus of the present study, were isolated from convalescent or vaccinated individuals, used different V genes (except for C13B8 and C13A7 that were clonally related) and displayed a high load of somatic mutations (7-14% in VH, 2-8% in VL at the nucleotide level, **Table S1**). These results illustrate the utility of a simple high-throughput method based on multiple parallel screening steps of memory B cells to isolate broadly reactive and even pan-reactive coronavirus mAbs.

## Pan-coronavirus mAbs bind to the fusion peptide and acquire affinity and breadth through somatic mutations

Using SARS-CoV-2 S_1_, RBD and S_2_ proteins, as well as 15-mer linear peptides covering the entire S sequence, the specificities of all seven pan-coronavirus mAbs were mapped to the K_811_PSKRSFIEDLLFNK_825_ sequence in the S_2_ subunit (**Fig. 2A**). This sequence spans the S_2_’ cleavage site (R815) and the fusion peptide N-terminal region, which is essential for membrane fusion (*28*) and is highly conserved amongst all genera of the *Orthocoronavirinae* subfamily, including the alpha, beta, gamma, and delta coronaviruses, as well as all SARS-CoV-2 variants sequenced to date (**Fig. S3, A** and **B**). Notably, the seven pan-coronavirus mAbs bound with different EC50 to the prefusion HexaPro and PentaPro S trimers as well as to prefusion and postfusion S_2_, likely due to the presence of the F817P mutation in HexaPro S (**Fig. 2A**).

**Fig. 2.**
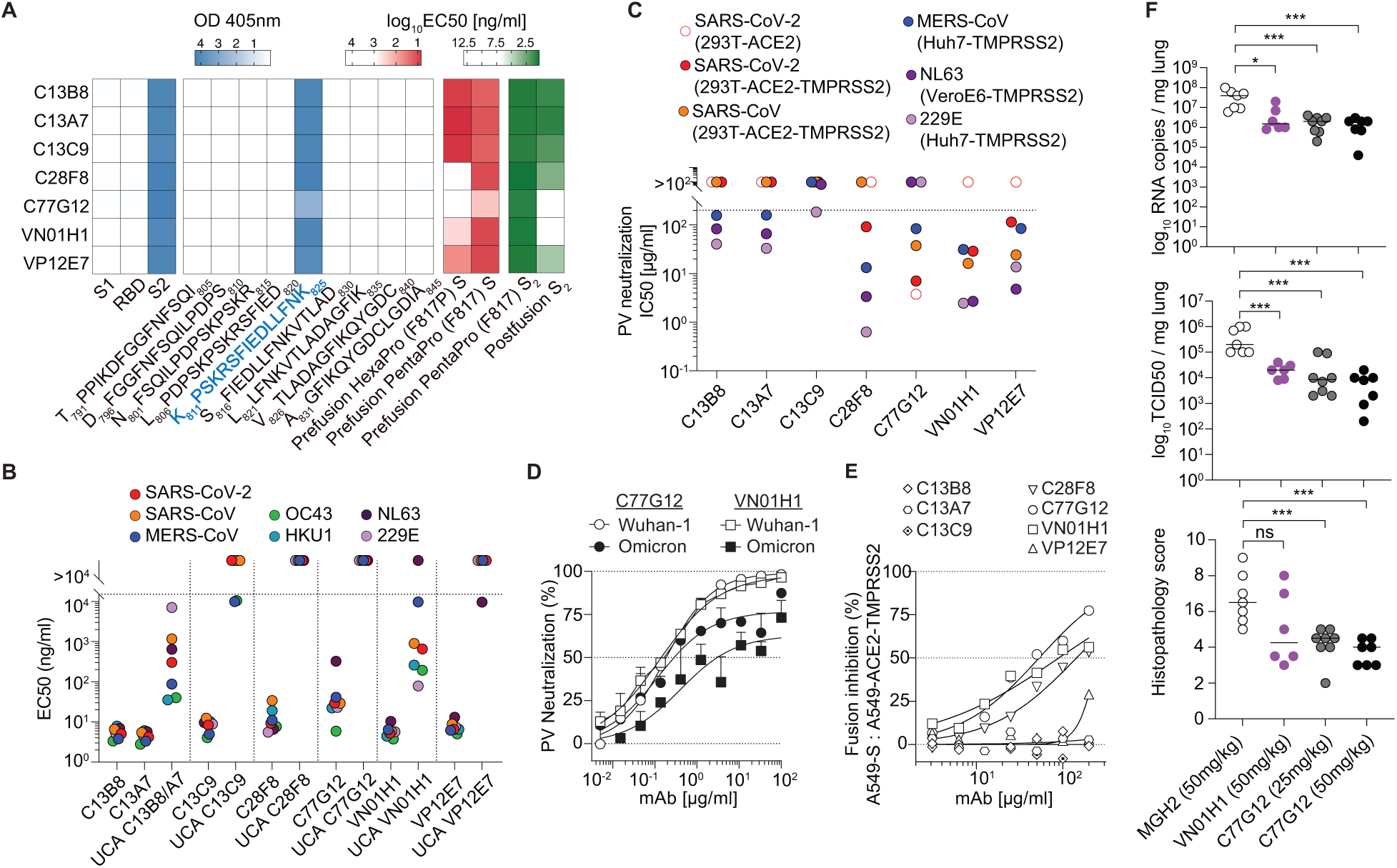
Pan-coronavirus mAbs target the fusion peptide and acquire affinity and breadth through somatic mutations. (**A**) Binding profiles of the seven pan-coronavirus mAbs. Epitope mapping was performed using SARS-CoV-2 protein fragments (S_1_, RBD, S_2_) and 15-mer overlapping synthetic peptides of the indicated sequences and measured by ELISA (OD405nm shown). The pan-coronavirus antibodies recognize the fusion peptide epitope highlighted in blue. EC50 values of the seven pan-coronavirus mAbs to prefusion HexaPro (F817P) S, prefusion PentaPro (F817) S, prefusion PentaPro (F817) S_2_ and postfusion S_2_ are also shown. (**B**) EC50 comparison of pan-coronavirus mAbs and their respective germline reverted unmutated common ancestors (UCAs) to different HCoV S proteins. C13B8 and C13A7 are sister clones and thus share a single UCA. Shown is one representative experiment out of at least two performed. (**C**) Pan-coronavirus mAbs were assessed for their ability to neutralize SARS-CoV-2, SARS-CoV, MERS-CoV, NL63 and 229E pseudoviruses in the indicated target cell lines. For each HCoV pseudotyped assay, all mAbs were compared in parallel. Scatter plot shows the collated IC50 neutralizing titers. Shown are data from one representative experiment out at least two performed. (**D**) VN01H1 and C77G12 mAbs were tested for their ability to neutralize SARS-CoV-2 Wuhan-Hu-1 and Omicron BA.1 variant pseudoviruses. (**E**) Pan-coronavirus mAbs were assessed for their ability to inhibit the fusion of SARS-CoV-2 S-expressing (A549-S) and ACE2-TMPRSS2-expressing (A549-ACE2-TMPRSS2) A549 cells. Inhibition of fusion values are normalized to the percentage of fusion without mAb (100%) and to that of fusion of wild-type A549 cells (0%). Shown are data from one representative experiment out two. (**F**) The prophylactic efficacy of VN01H1 (50 mg/kg), C77G12 (25 mg/kg and 50 mg/kg) and negative control mAb MGH2 (50 mg/kg) were tested in hamsters challenged with SARS-CoV-2 P.1 (Gamma) VOC. Viral RNA loads (top), replicating virus titers (middle), and histopathological scores (bottom) are shown. *p < 0.05, ***p < 0.001, Mann-Whitney test.

To explore the ontogeny of fusion peptide-specific pan-coronavirus mAbs, we compared the binding properties of mature mAbs to their unmutated common ancestors (UCAs) (**Fig. 2B** and **Fig. S2B**). The UCAs of C13B8, C13A7 (the two clonally related mAbs) and VN01H1 already exhibited broad reactivity to HCoV S proteins but had lower affinity. In contrast, the C13C9 UCA bound only to the beta HCoVs OC43 and MERS-CoV S, whereas the VP12E7 UCA bound only to the alpha HCoV NL63 S, with low avidity in both cases. Finally, the UCAs of C28F8 and C77G12 did not bind to any HCoV S proteins tested. Given the lack of a common V gene usage, these findings suggest that pan-reactive fusion peptide-specific mAbs can mature through multiple pathways and acquire high affinity and cross-reactivity through somatic mutations, possibly as a consequence of priming by endemic coronavirus infection followed by SARS-CoV-2 boost.

Considering the conservation of the fusion peptide region, we next asked if pan- or broad reactivity would be a property shared by most fusion peptide-specific antibodies. IgGs secreted upon polyclonal activation of memory B cells from 71 convalescent individuals were screened for binding to a pool of peptides comprising the fusion peptide sequences of alpha and beta HCoVs, as well as to SARS-CoV-2 S protein. Cultures producing fusion peptide-specific antibodies were detected at low frequency and only in 19 individuals (**Fig. S4A**). Although nearly all fusion peptide-reactive antibodies bound to SARS-CoV-2 S protein, only 9 out of 30 were pan-reactive, while the remaining antibodies showed different degrees of cross-reactivity (**Fig. S4B**). Thus, although the fusion peptide region in the coronavirus S protein is immunogenic (*29–33*) and conserved, pan-reactivity is the property of a minority of fusion peptide-specific mAbs.

## Anti-fusion peptide mAbs show varied neutralizing activity and can reduce viral burden in vivo

We next tested the neutralizing activity of all seven mAbs against alpha and beta HCoV pseudoviruses in TMPRSS2-expressing target cell lines. Despite binding to the same motif, these mAbs displayed heterogeneous neutralizing potencies. Most notably, VN01H1 and VP12E7 neutralized all pseudotyped alpha and beta HCoVs tested (SARS-CoV-2, SARS-CoV, MERS-CoV, NL63 and 229E), as well as pre-emergent bat *sarbecovirus* WIV-1 (**Fig. 2C** and **Fig. S5, A** and **B**). In addition, C77G12 neutralized all beta coronaviruses and showed efficient neutralization of SARS-CoV-2 Wuhan-1 and Omicron BA.1 and inhibition of cell-cell fusion (**Fig. 2, D** and **E**). These results suggest that fusion peptide-specific mAbs can impede S proteolytic activation or fusogenic rearrangements, thereby inhibiting membrane fusion.

In view of the broad reactivity of VN01H1 and the relatively high potency of C77G12, we further assessed the protective efficacy of these mAbs in vivo. In the Syrian hamster model of SARS-CoV-2 P.1 (Gamma) infection, prophylactic administration of either mAb at high doses reduced RNA copies and lung viral titers, and ameliorated lung pathology, in a statistically significant manner (**Fig. 2F**). In summary, these findings demonstrate that fusion peptide-specific mAbs display protective efficacy in vivo, albeit with moderate potency.

## Structural basis for pan-coronavirus mAb cross-reactivity

To gain insights into the epitope recognized by fusion peptide-specific pan-coronavirus mAbs, we first performed substitution scan analysis on the six clonally unrelated mAbs. All mAbs displayed a core binding motif at I_818_EDLLFNK_825_ (**Fig. S6A**) but C28F8, C77G12, VN01H1, and VP12E7, showed a 3-amino acid expanded footprint spanning the N-terminal R_815_SF_817_ residues, comprising the S_2_’ cleavage site at position R_815_.

We then determined crystal structures of five mAbs (C13B8, C13C9, C77G12, VN01H1 and VP12E7) in complex with the K_811_PSKRSFIEDLLFNK_825_ fusion peptide at 2.1 Å, 2.1 Å, 1.7 Å, 1.86 Å and 2.5 Å, resolution, respectively (**Fig. 3** and **Fig. S6, B** to **D** and **Table S2**). All five mAbs bind to overlapping epitopes in the fusion peptide through interactions involving the heavy and light chains. The strict conservation or the conservative substitution of key residues involved in mAb recognition (R_815_, S_816_, I_818,_ E_819_, D_820_, L_821_, L_822_, F_823_, N_824_ and K_825_) across the *Orthocoronavirinae* subfamily explains the unprecedented cross-reactivity of these fusion peptide-specific mAbs (**Fig. S3** and **Table S4**). The overall architecture of the fusion peptide in the C13C9- bound complex structure is most similar to that observed in the prefusion S trimer (PDB 6VXX) (*3, 34*) (**Fig. 3C** and **Fig. S6B**). The SARS-CoV-2 S fusion peptide adopts a similar conformation in the two structures determined in complex with VN01H1 or VP12E7, which is distinct from the conformation observed in prefusion SARS-CoV-2 S trimeric structures (*3, 34*) (**Fig. 3, B** and **C** and **Fig. S6C**). Specifically, residues _813_SKR_815_ refold from an extended conformation in prefusion S to an ɑ-helical conformation in the two Fab-bound peptide structures, thereby extending the ɑ-helix found at the N-terminal region of the fusion peptide. The fusion peptide residues P_812_-R_815_ adopt an extended conformation, distinct from prefusion S, in the C77G12-bound complex structure (**Fig. 3, A** and **C**), whereas these residues are disordered in the C13B8-bound structure (**Fig. 3C** and **Fig. S6D**). The conserved residue R815, which is the S_2_’ site of proteolytic processing upon receptor binding for membrane fusion activation, is engaged in electrostatic interactions with the C13C9, VN01H1, VP12E7 and C77G12 Fabs and therefore buried at the interface with their paratopes (**Table S4**). Since pre-incubation of a soluble native-like SARS-CoV-2 S ectodomain trimer with fusion peptide-specific mAbs did not prevent S2X58-induced triggering of fusogenic conformational changes (*35*) (**Fig. S7**), these mAbs likely inhibit TMPRSS2 cleavage of the S_2_’ site (via steric hindrance) and in turn activation of membrane fusion. Although residue F817 is not part of the epitope (**Table S4**), the F817P substitution present in the ‘HexaPro’ construct likely prevents the adoption of the extended ɑ-helical conformation observed in the structures bound to VN01H1 or VP12E7 (due to restricted backbone torsion angles), resulting in dampened binding in ELISA (**Fig. 2A**).

**Fig. 3.**
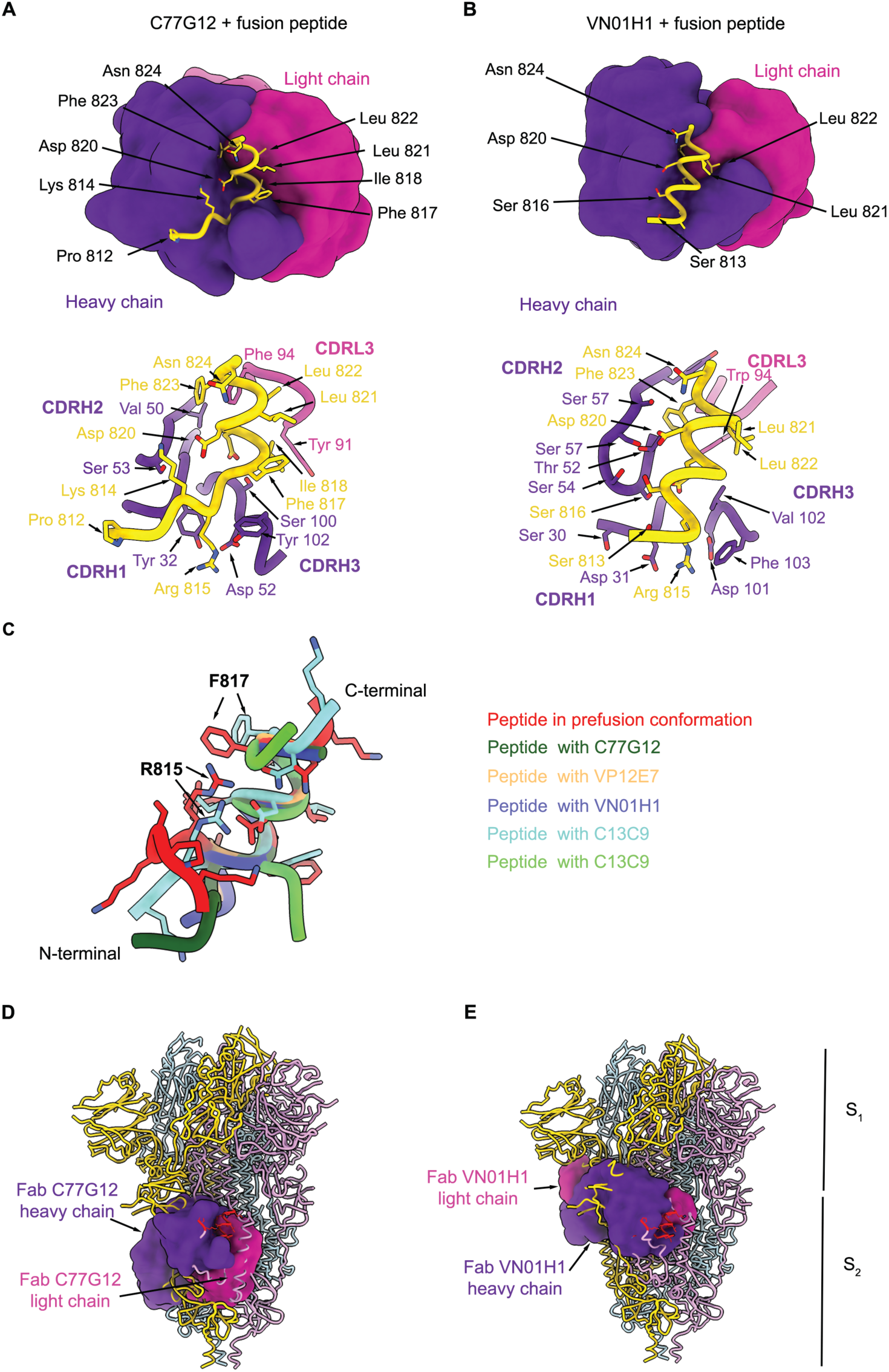
Fusion peptide antibodies target a cryptic epitope. (**A** and **B**) (Top) Surface representation of the crystal structures of the C77G12 (**A**) and VN01H1 (**B**) Fabs in complex with SARS-CoV-2 fusion peptide epitope. (Bottom) Ribbon representation of the corresponding structures highlighting the interactions of Fab heavy and light chain CDRs with the fusion peptide (selected regions are shown for clarity). (**C**), Ribbon representation of the fusion peptides in the Fab-bound complexes superimposed with the fusion peptide in prefusion SARS-CoV-2 S (PDB 6VXX). (**D** and **E**) Superimposition of the C77G12-bound (**D**) or VN01H1-bound (**E**) fusion peptide structures to prefusion SARS-CoV-2 S uncovering the cryptic nature of the epitope. The Fabs are shown as surfaces whereas S is rendered as ribbons. Each SARS-CoV-2 S protomer is colored distinctly (light blue, pink, and gold) and Fab heavy and light chains are colored purple and magenta, respectively.

Superimposition of the mAb-fusion peptide complexes with available prefusion S structures (PDB 6VXX) (*3, 36*) revealed that the targeted epitope is buried toward the core of the S trimer and is therefore inaccessible (**Fig. 3, D** and **E** and **Fig. S6, B** to **D**), likely explaining the lack of detectable complexes of Fabs with prefusion S trimers during single particle electron microscopy analysis. Taken together, these findings suggest that the epitope recognized by fusion peptide-specific mAbs is cryptic and may become accessible only transiently (*37*).

## The fusion peptide epitope is unmasked by ACE2 binding

To investigate how fusion peptide-specific mAbs might bind to native S trimers and exert neutralizing activity, we transfected 293T cells to express coronavirus S proteins on the cell surface (**Fig. 4A** and **Fig. S8**). Interestingly, all fusion peptide-specific mAbs showed only marginal binding to SARS-CoV-2 S-expressing 293T cells, as compared to control mAbs targeting the RBD (C94) or the stem helix (C21E3). Remarkably, addition of soluble ACE2 enhanced the binding of all fusion peptide antibodies to native SARS-CoV-2 S protein to levels comparable to that of control mAbs, indicating that receptor engagement induces a conformational change that exposes the cryptic fusion peptide epitope (**Fig. 4A**). Importantly, this ACE2-dependent enhancement of binding was not observed in 293T cells expressing SARS-CoV-2 S protein harboring the 2P (K986P, V987P) prefusion-stabilizing mutations (*3, 36*) that lay outside of the epitope (**Fig. 4A** and **Fig. S8**), suggesting an impediment of the receptor-induced allosteric conformational changes. Enhanced binding of fusion peptide-specific mAbs was also observed in SARS-CoV and MERS-CoV S expressing 293T cells in the presence of the corresponding receptors ACE2 (*38*) and DPP4 (*39*). In contrast, these mAbs bound efficiently to NL63 and 229E S 293T cells, independently of receptor engagement by ACE2 (*40*) or APN (*41*), respectively (**Fig. 4A** and **Fig. S8**).

**Fig 4.**
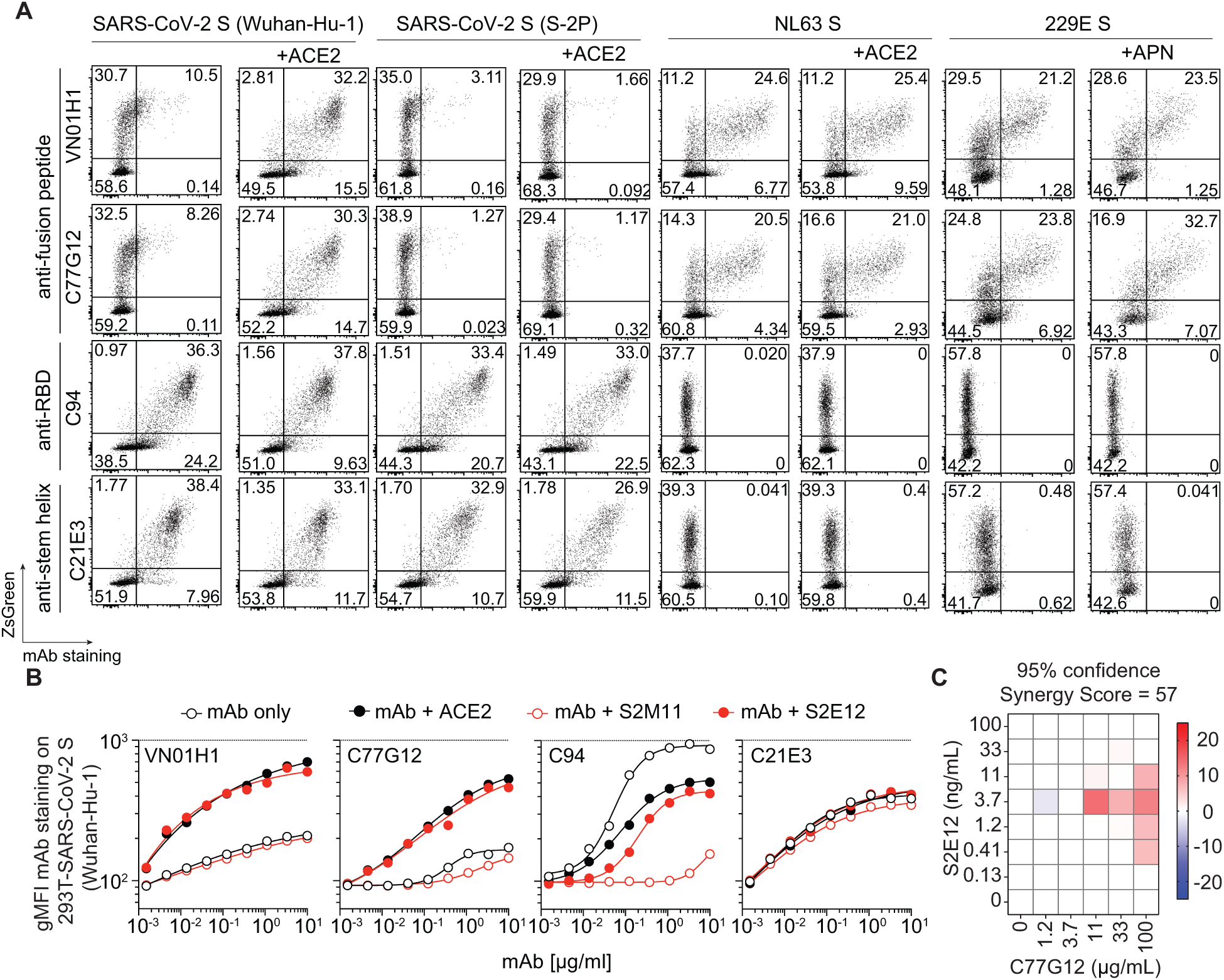
The SARS-CoV-2 fusion peptide is unmasked following ACE2 receptor engagement. (**A**) Binding of fusion peptide-specific mAbs VN01H1 and C77G12 (8 μg/ml), as well as RBD-specific C94 and stem helix-specific C21E3 mAbs (8 μg/ml) on 293T cells transiently co-transfected with plasmids encoding ZsGreen and SARS-CoV-2 S Wuhan-Hu-1, SARS-CoV-2 S-2P, NL63 S or 229E S, in the presence or absence of receptor ACE2 or APN, as measured by flow cytometry. (**B**) Titrating doses of fluorescently labelled antibodies were co-incubated with 293T-S cells for 2 h at room temperature in the presence or absence of recombinant ACE2 fusion protein (27 µg/ml), ACE2-mimicking mAb S2E12 (20 µg/ml) or S2M11 (20 µg/ml), a mAb that locks S trimer in a closed conformation. The line in the scatter plot is a reference for the maximum gMFI of staining with control anti-RBD C94 mAb. Shown is one representative experiment out of two. (C) Synergy experiment was performed against SARS-CoV-2 (D614G) on Vero E6-TMPRSS2 at indicated concentrations of S2E12 and C77G12 mAbs alone or in combination. Shown is the synergy map where synergy score is calculated using synergy-scoring model (MacSynergy II) (*61*).

It was previously shown that certain mAbs can mimic ACE2 binding and trigger fusogenic activity (*42, 43*). Indeed, the addition of S2E12, an ACE2-mimicking mAb, but not S2M11, a mAb which locks SARS-CoV-2 S trimer in the closed state (42), was also able to enhance binding of the fusion peptide antibody (**Fig. 4B**). Consistent with this finding, we observed a synergy in pseudovirus neutralization when C77G12 was used in combination with S2E12, suggesting that fusion-peptide antibodies may be more effective in the context of a polyclonal response against the RBD. Collectively, these findings indicate that the fusion peptide epitope only becomes accessible upon receptor-induced S conformational changes in *sarbecovirus* SARS-CoV-2, SARS-CoV and *merbecovirus* MERS-CoV, whereas it is more readily accessible in the *alphacoronavirus* 229E and NL63, possibly due to molecular breathing or structural flexibility.

## Discussion

By screening a large number of memory B cells from immune donors using a panel of S proteins, we identified a new class of mAbs that target the fusion peptide region, some of which have unprecedented breadth being able to bind and neutralize both alpha and beta coronaviruses. Structural analysis demonstrates that despite targeting a conserved 15 amino acid motif, these mAbs use different V genes and exhibit different binding modalities. The finding that the UCAs of these antibodies bind preferentially to common cold coronaviruses and that the antibodies acquire affinity and breadth through somatic mutations suggest that their elicitation require multiple rounds of selection, possibly by heterologous stimulation by different coronaviruses. A complex developmental pathway has been reported for HIV-1 neutralizing antibodies and may be a general requirement for antibodies that recognize highly constrained epitopes (*44, 45*).

Previous studies identified serum antibodies to the fusion peptide of SARS-CoV-2 and showed, through depletion or peptide inhibition experiments, that such antibodies can contribute to the serum neutralizing activity in a polyclonal setting (*29–33*). We show here that some fusion peptide-specific mAbs have direct neutralizing activity on pseudoviruses in vitro and reduce viral burden in vivo when administered at high doses in a hamster challenge model. Although the neutralizing activity of these mAbs is low when used alone, it is possible that in the context of a polyclonal response they may synergize with other antibodies that favor the exposure of the fusion peptide region, as shown here with the S2E12 mAb.

Although the conformational rearrangements of the fusion peptide region enabling S_2_’ cleavage have thus far eluded structural definition, several prior studies described the biochemical events of S_2_’ proteolytic processing for SARS-CoV-2 (*46, 47*) and MERS-CoV (*48, 49*). The mAbs isolated in this study provided a tool for defining an allosteric conformational change in the S protein that is triggered by receptor binding or by receptor-mimicking antibodies (*42, 43*). This conformational change, which is abolished by introduction of stabilizing 2P mutations at a distant site, may be required to expose the fusion peptide region to the proteolytic cleavage by TMPRSS2 or endosomal cathepsins, resulting in the activation of the S protein fusogenic activity (*28, 50*). The finding that fusion peptide epitopes are readily accessible in alphacoronaviruses even in the absence of receptor engagement is consistent with the higher neutralizing activity of fusion peptide-specific mAbs against NL63 and 229E and suggests that such conformational change might occur spontaneously as a consequence of molecular breathing. The inclusion of the prefusion stabilizing 2P mutations implies that current mRNA vaccines may disfavor the elicitation of an antibody response to the fusion peptide region.

The fusion peptide region is highly conserved across the *Orthocoronavirinae* subfamily and, in view of its functional relevance, may be less prone to viral escape due to the potential fitness loss of mutants (*51, 52*). Broadly neutralizing mAbs to the fusion peptide region of influenza hemagglutinin and HIV-1 gp120 have guided the design of universal vaccines against these highly variable pathogens (*53–59*). Similarly, the pan-coronavirus mAbs described in this study may be used as probes for the design of immunogens that can better unmask the fusion peptide region and elicit a broadly protective antibody response. Interestingly, the same region was also found to stimulate broadly reactive CD4 T cells, providing a cue for intramolecular help in the generation of such antibodies (*60*).

## Acknowledgements

We thank all study participants who donated blood and devoted time to our research. We thank Maira Biggiogero, Alessandra Franzetti Pellanda, Elena Picciocchi, Tatiana Terrot, Sonia Tettamanti, Tiziana Urbani, Luisa Vicari and all personnel at the hospitals and nursing homes for providing blood samples, Daniela Vaqueirinho, Sandra Jovic, Isabella Giacchetto Sasselli, Rahel Schmidt and Xinlei Xi from the Sallusto laboratory for their help with blood processing, Manfred Kopf (ETH Zurich) for providing the pLVX-puro-ACE2 transfer plasmid, Hideki Tani (University of Toyama) for providing the reagents necessary for preparing VSV pseudotyped viruses, and Marc Weisshaar (ETH Zurich) for his artistic rendition of Fig. S1.

## Funding

The study was in part funded by:

Henry Krenter Foundation (FS)

ERC AdG n. 885539 ENGRAB (AL)

National Institute of Allergy and Infectious Diseases DP1AI158186 and HHSN272201700059C (DV)

Pew Biomedical Scholars Award (DV)

Investigators in the Pathogenesis of Infectious Disease Awards from the Burroughs Wellcome Fund (DV)

Fast Grants (DV)

Natural Sciences and Engineering Research Council of Canada (MM)

University of Washington Arnold and Mabel Beckman cryoEM center

National Institute of Health grant S10OD032290 (DV)

Beamline 5.0.1 at the Advanced Light Source at Lawrence Berkley National Laboratory EOC research funds

DV is an Investigator of the Howard Hughes Medical Institute

FS and the Institute for Research in Biomedicine are supported by the Helmut Horten Foundation

## Competing interests

JSL, JJ, FS, AL, ACa are currently listed as an inventor on multiple patent applications, which disclose the subject matter described in this manuscript. AL, DC, FS, JN, MM-R, LAP and DP may hold shares in Vir Biotechnology. The Veesler laboratory and the Sallusto laboratory have received sponsored research agreements from Vir Biotechnology Inc. The other authors declare no competing interests.

## Authors contribution

Conceptualization: JSL, MAT, AL, DV, FS

Methodology: JSL, JJ, MAT, MM, DP, AC, SB, JEB, AJ, ACW, DM, FM, PP, DJ, MF, RA, BW, JNo

Investigation: JSL, JJ, MAT, MM, DP, AC, SB, JEB, AJ, ACW, DM, FM, PP, DJ, MF, RA, BW, JNo, JNe, MM, LAP, CG, PF, ACe

Funding acquisition: FS, DV, AL

Supervision: JSL, DC, DV, AL, FS

Writing, review & editing: JSL, MAT, DC, DV, AL, FS

## Data and materials availability

All data associated with this manuscript are available in the main text or the supplementary materials, including FACS data gating strategy. The crystal structures will be deposited to the protein data bank (PDB). All further relevant source data that support the findings of this study are available from the corresponding authors upon reasonable request. Materials are available through materials transfer agreements (MTAs).

---

**Fig. S1.**
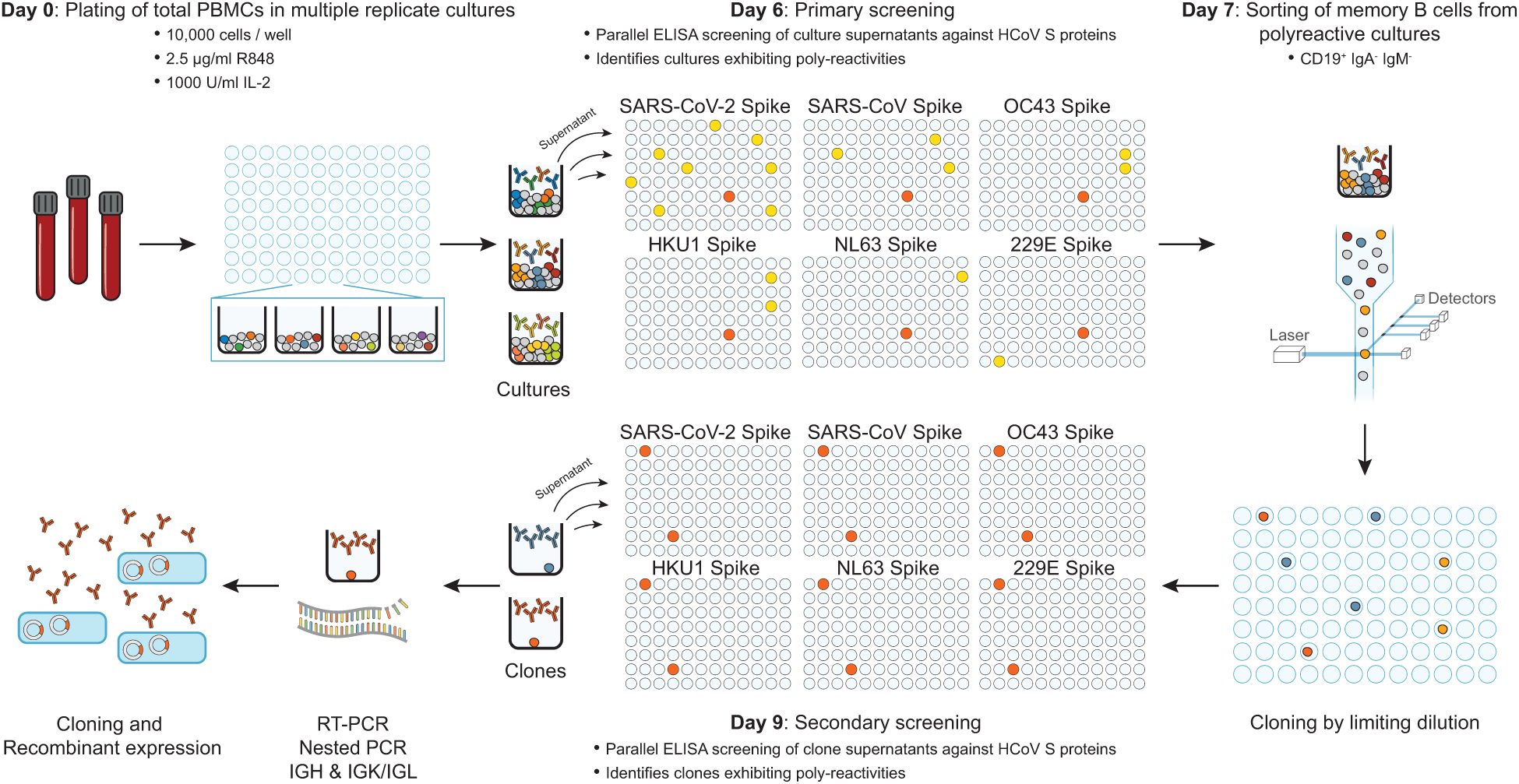
An improved method that combines the high-throughput screening approach and limiting dilution cloning to isolate B cells of unique characteristics. Total PBMCs isolated from Ficoll density centrifugation were plated in replicate cultures at densities of 10^4^ cells per well in the presence of 2.5 µg/ml of TLR agonist R848 and 1,000 U/ml IL-2. Six days later, the specificities of the secreted IgG antibodies in the culture supernatants from each culture were screened against different HCoV S antigens in parallel (primary screening). Cultures that exhibit cross-reactive binding patterns (shown as red well) were next isolated as CD19^+^ IgM^−^ and IgA^−^ to enrich for IgG-secreting memory B cell blasts (*62*) and cloned by limiting dilution at 0.7 cell/well. Two days post cloning, the culture supernatants of the clones underwent secondary screening with the same panel of antigens to validate their binding profiles observed during primary screening. Clones which exhibit the same binding profiles as primary screening were selected for BCR retrieval by reverse transcription followed by nested PCR reactions. Paired IGH and IGK/IGL were cloned into expression vector and transfected into Expi293 cells for recombinant antibody expression. Recombinantly expressed antibodies were tested for their specificities to the S antigens as validation.

**Fig. S2.**
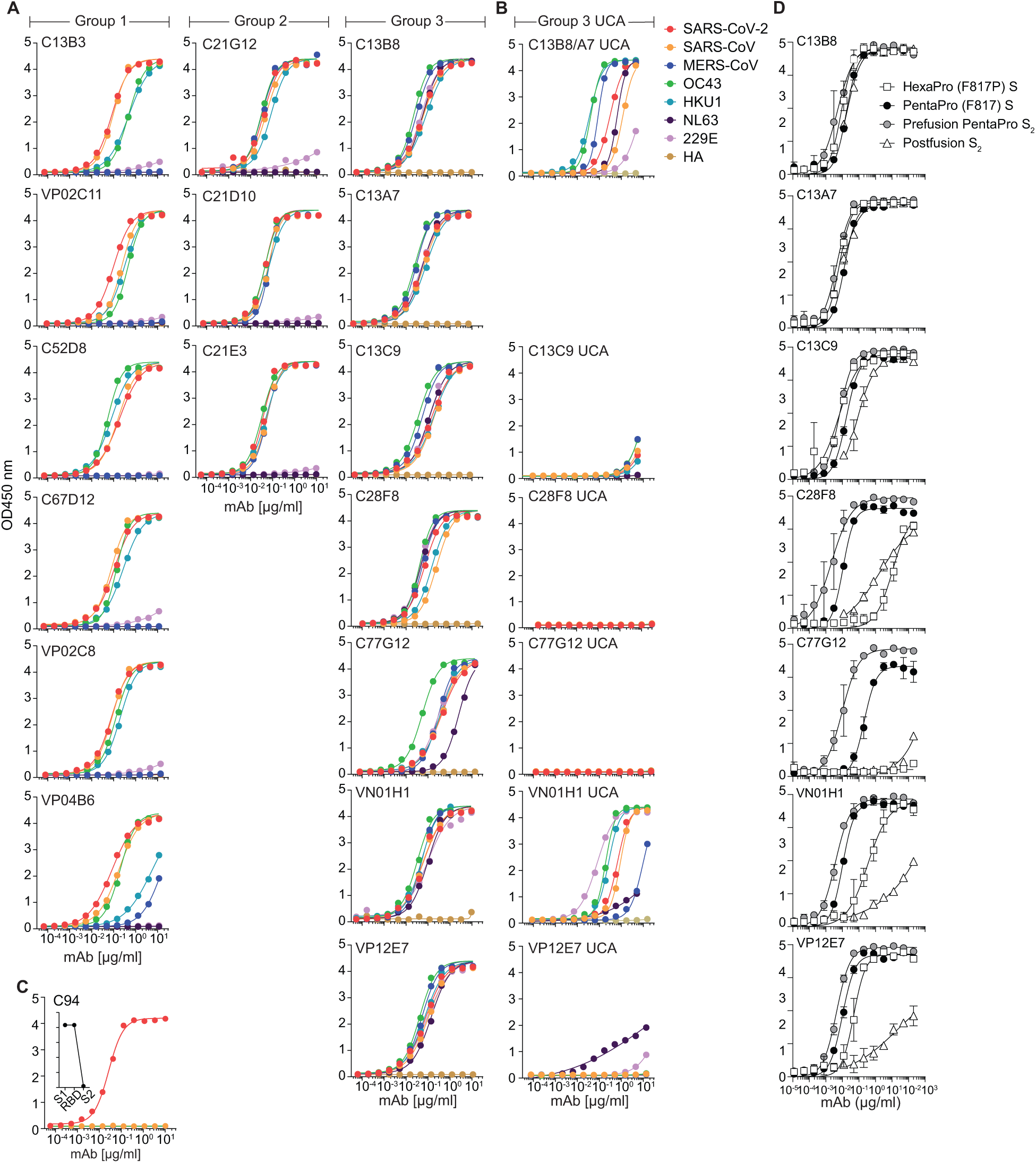
The binding profiles of all 16 HCoV-cross-reactive mAbs. (**A** and **B**) The binding profiles of all 16 cross-reactive mAbs (**A**) and respective unmutated common ancestors (UCAs) of Group 3 mAbs (**B**) were tested against plate-immobilized HCoV S protein antigens, as well as control antigen H1N1 haemagglutinin (HA) by ELISA. (**C**) Shown is the binding profile of the anti-RBD mAb C94, used as a control. Inset represents the domain mapping (S_1_, RBD, S_2_) of mAb C94 where the relative maximum OD 405 nm value is shown. Shown are data from one representative experiment out of at least two performed. (**D**) Bindings of all seven anti-fusion peptide mAbs to prefusion SARS-CoV-2 S S HexaPro (F817P), SARS-CoV-2 S PentaPro (F817), SARS-CoV-2 S_2_ PentaPro in prefusion conformation and SARS-CoV-2 S_2_ in postfusion conformation were analyzed by ELISA. Shown is one representative experiment out of two.

**Fig. S3.**
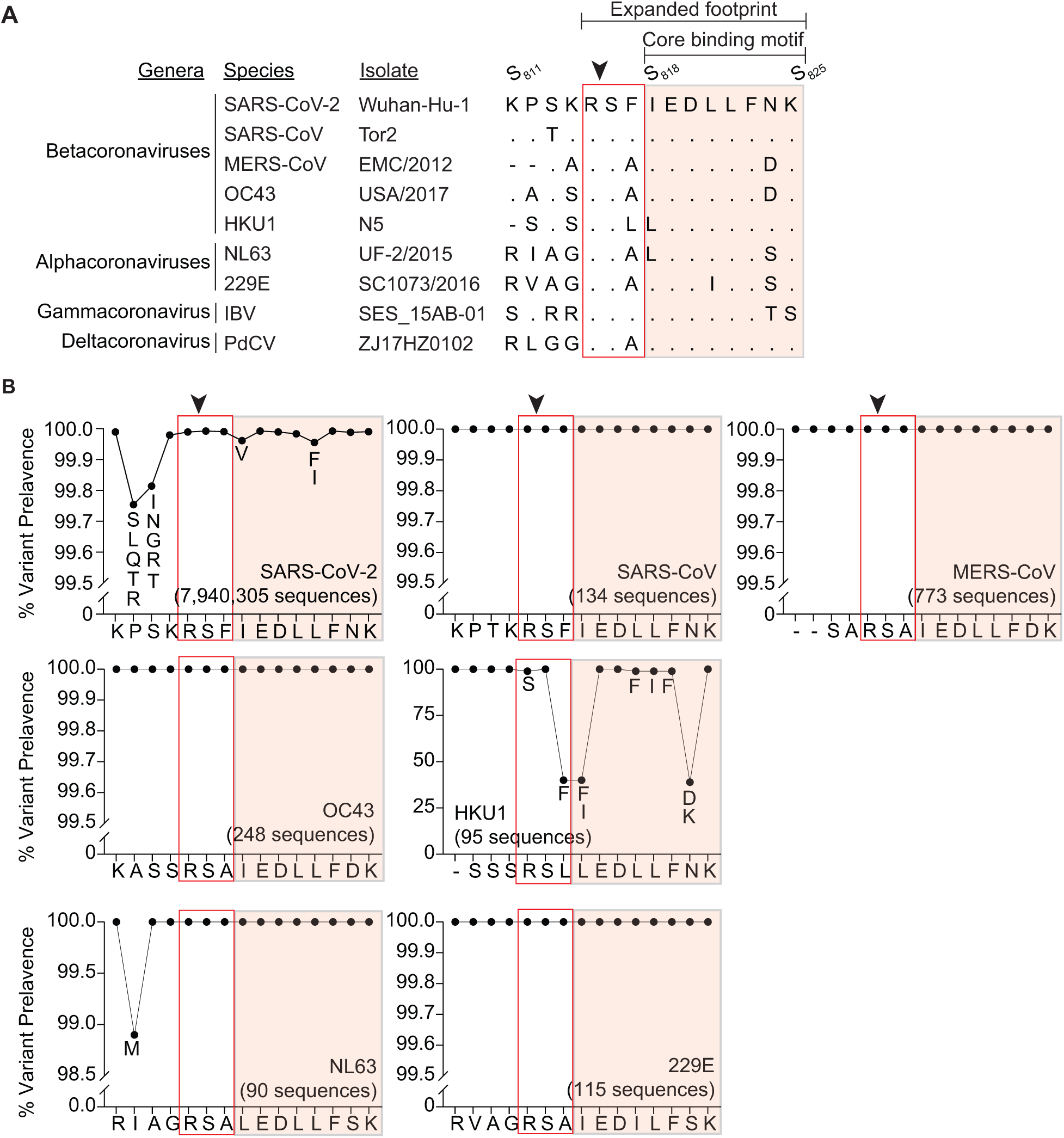
The fusion peptide epitope is highly conserved amongst *Orthocoronavirinae* subfamily and mutations in this region in circulating SARS-CoV-2 variants are uncommon. (A) Alignment of the fusion peptide motif of alpha, beta, gamma and delta coronaviruses. (**B**) Frequency of mutations in the fusion peptide region is analyzed based on deposited HCoV sequences at GISAID (SARS-CoV-2) and NCBI virus database (all others), as of 9^th^ February 2022. SARS-CoV-2 (7,940,305 sequences), SARS-CoV (134 sequences), MERS-CoV (773 sequences), OC43 (248 sequences), HKU1 (95 sequences), NL63 (90 sequences), 229E (115 sequences). Arrows indicate S_2_’ cleavage site. Core binding motif and expanded footprint identified from Fig. S6A are shown.

**Fig. S4.**
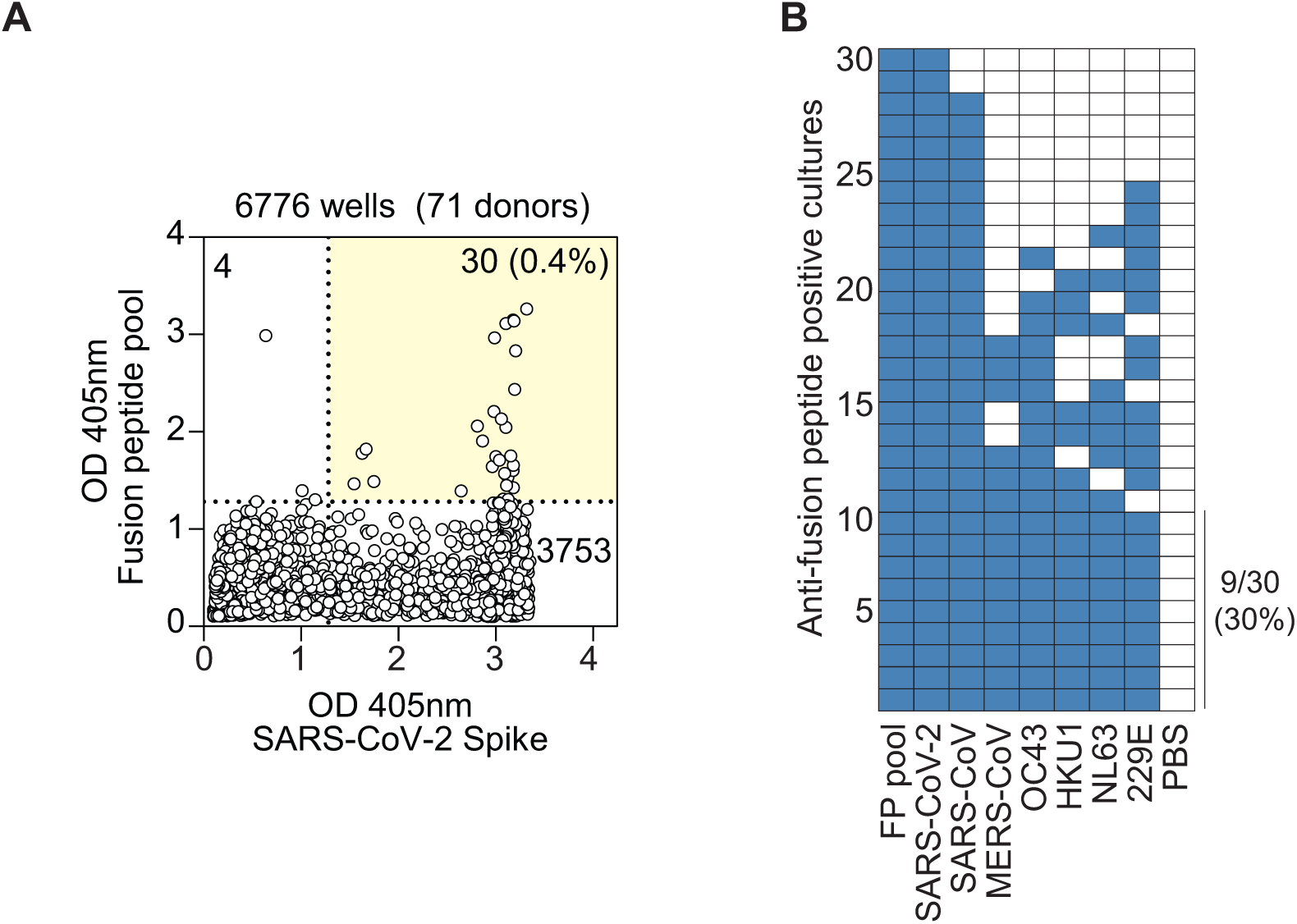
Fusion peptide reactive antibodies are rare and not all fusion peptide antibodies are pan-reactive. (**A)** Total PBMCs from 71 COVID-19 convalescent individuals were plated in replicate 96 wells (3 × 10^4^ cells/well) and stimulated with R848 (2.5 µg/ml) and IL-2 (1,000 U/ml). Six days later, the supernatant of each culture was screened, in parallel, for the specificities of the secreted antibodies to a pool of 15-mer synthetic fusion peptides from SARS-CoV-2, SARS-CoV, MERS-CoV, OC43, HKU1, NL63 and 229E, as well as to SARS-CoV-2 S protein by ELISA. Each circle in the scatter plot represents one culture and its respective OD 405nm values to fusion peptide pool and to SARS-CoV-2 S protein. (**B**) Cultures exhibiting reactivities to both fusion peptide pool and SARS-CoV-2 S protein (yellow quadrant in (**A**)) were further screened for their reactivities to other HCoV S proteins. Each row represents a fusion peptide- and SARS-CoV-2 S-double positive culture and each column shows the reactivities to the indicated S protein antigens. If OD 405nm value exceeds the cut-off value determined by average OD 405nm of PBS wells + 4*standard deviation, the culture was considered reactive to the antigen as indicated by a colored cell.

**Fig. S5.**
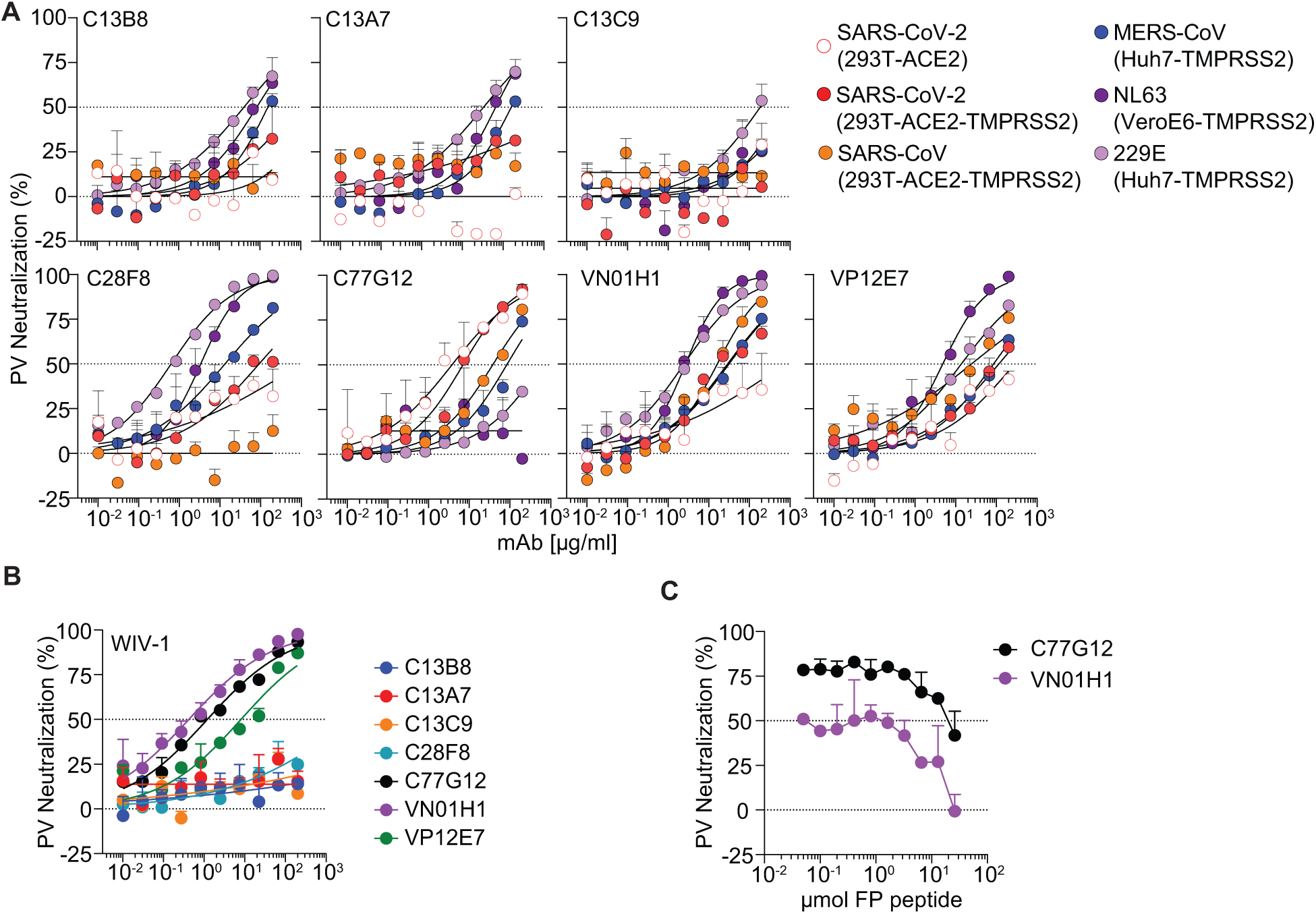
Anti-fusion peptide antibodies can neutralize human and animal alpha and beta coronaviruses. (**A**) Titrating doses of pan-reactive mAbs were assessed for their ability to neutralize: SARS-CoV-2, SARS-CoV, MERS-CoV, NL63 and 229E pseudoviruses in the indicated target cell lines. For each HCoV pseudotyped assay, all mAbs were compared in parallel. (**B**) Titrating doses of pan-reactive mAbs were assessed for their ability to neutralize WIV-1 in 293T-ACE2-TMPRSS2 cell line. (**C**) SARS-CoV-2 neutralization activity of VN01H1 and C77G12 (200 µg/ml) were competed with titrating doses of soluble fusion peptide.

**Fig. S6.**
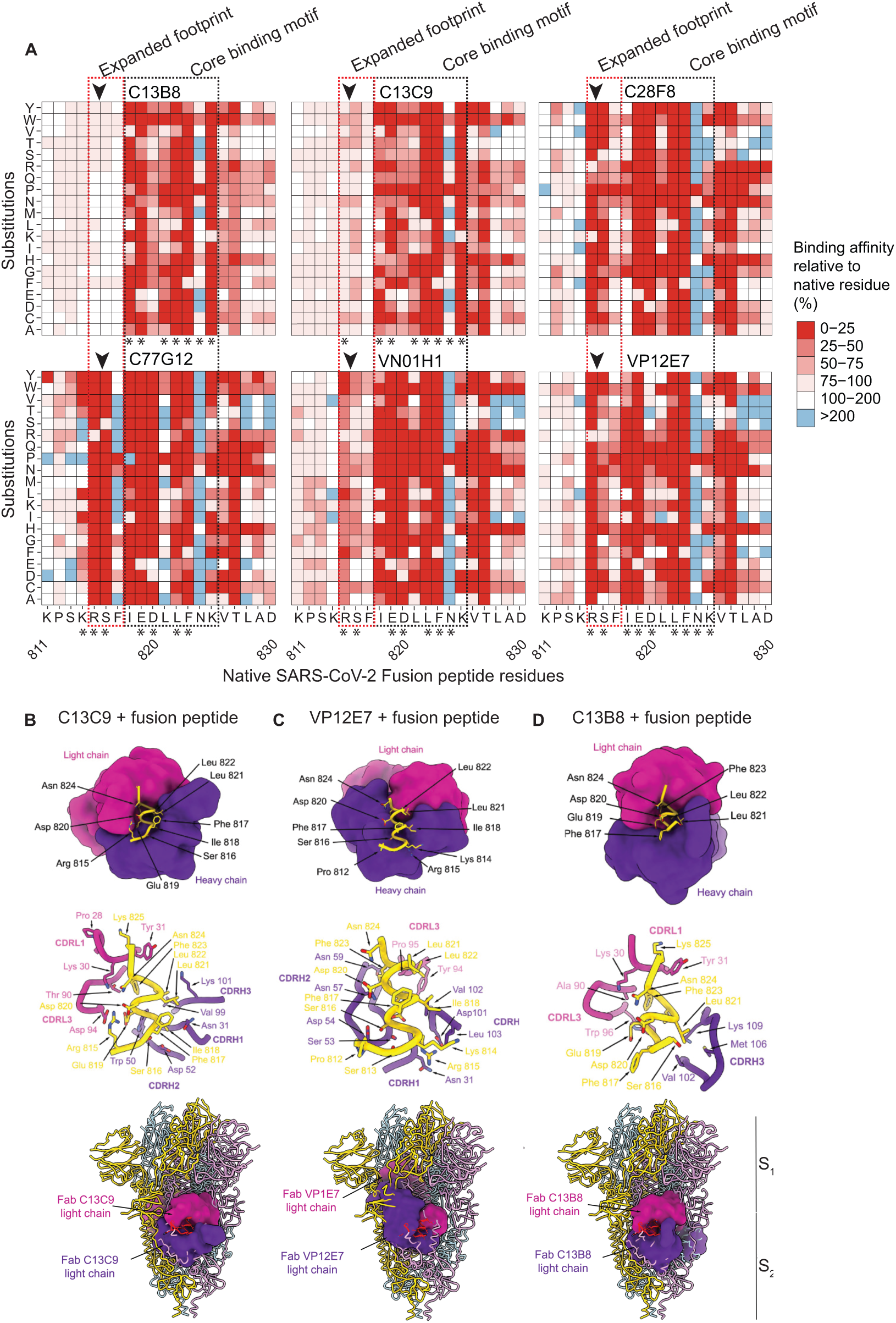
Varied binding profiles and the cryptic nature of anti-fusion peptide antibodies. (**A**) Heatmaps displaying the substitution scan analysis of six indicated mAbs (C13A7 and C13B8 are clonally related). Each amino acid in the fusion peptide epitope (*x*-axes) was substituted, step-wise, with all amino acids (*y*-axes), and the binding affinity of the antibody to each peptide variant was measured. Legend shows the binding affinity relative to the native residue. Epitope residues for each mAb defined based on obtained crystal structures are highlighted with asterisks (*). Arrows indicate S_2_’ cleavage site. Identified core binding motif and expanded footprint are shown. (**B** to **D**) Crystal structures of the C13C9 (**B**), VP12E7 (**C**) and C13B8 (**D**) Fabs (surface representation) in complex with SARS-CoV-2 fusion peptide epitope in ribbon representation (top panels). Ribbon representation of the crystal structures of Fab-bound complexes highlighting the interactions with the CDRs of the Fab heavy and light chains; only selected regions are shown for clarity (middle panels). Alignment between the fusion peptide in SARS-CoV-2 S in prefusion conformation (PDB 6VXX) with the fusion peptide (both in ribbon representation) in the crystal structure of the Fab-bound complex with C13C9, VP12E7 and C13B8 Fabs (surface representation) uncovering the cryptic nature of the epitope (bottom panels). Each SARS-CoV-2 S protomer is colored distinctly (light blue, pink, and gold). Fab heavy and light chains are colored purple and magenta, respectively.

**Fig. S7.**
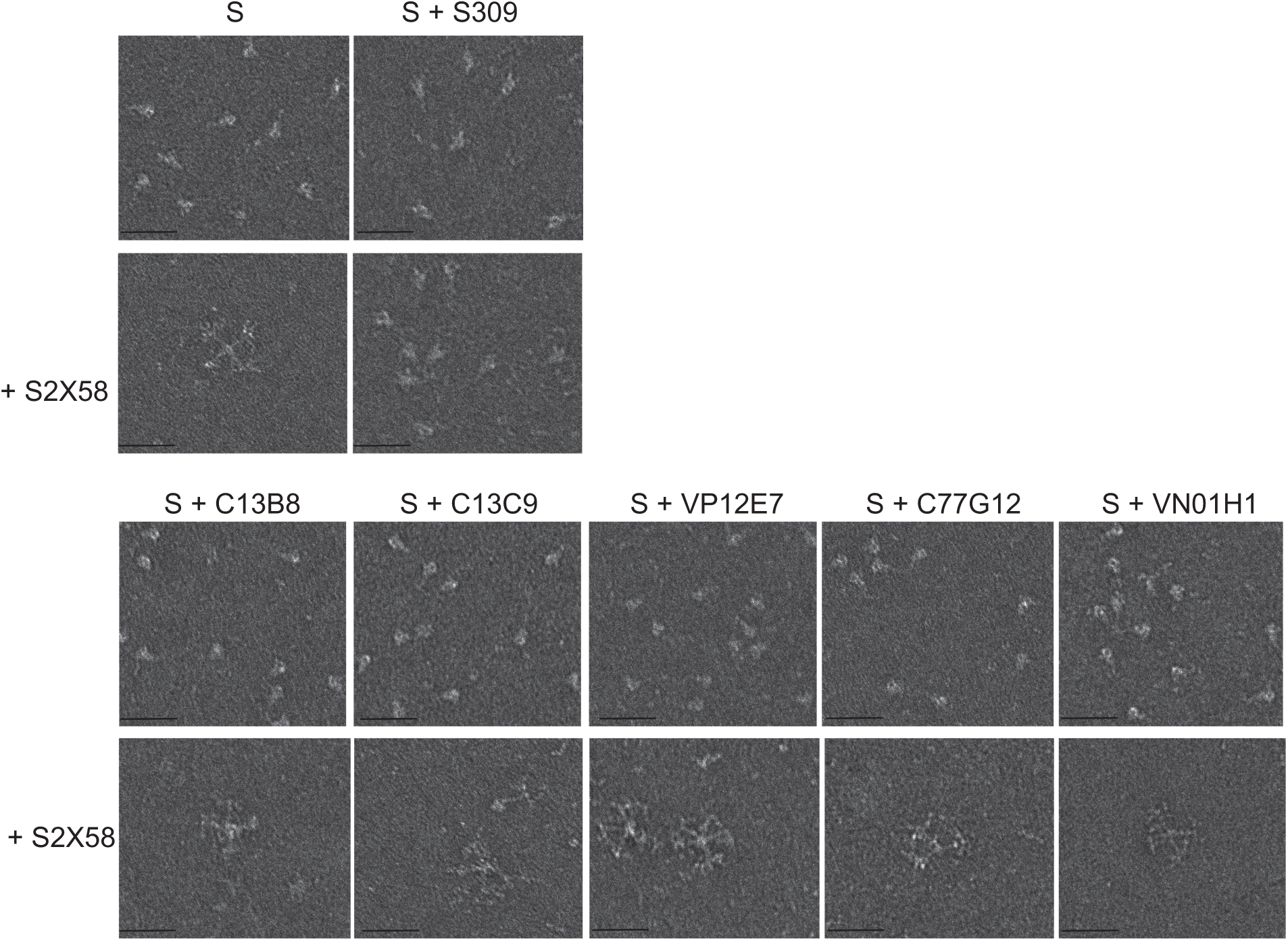
Fusion peptide-specific Fabs do not block fusogenic rearrangement of the S protein. Fusion peptide-specific Fabs were incubated for 1 h with native-like soluble prefusion S trimer of SARS-CoV-2 prior addition of the Fab S2X58 to induce the fusogenic rearrangement of soluble S trimers visible as rosettes by negative stain EM (scale bar: 50 nm). 30 micrographs per sample were analyzed. Negative control: S protein only incubated with S2X58. Positive control: S protein preincubated with Fab S309 before addition of S2X58.

**Fig. S8.**
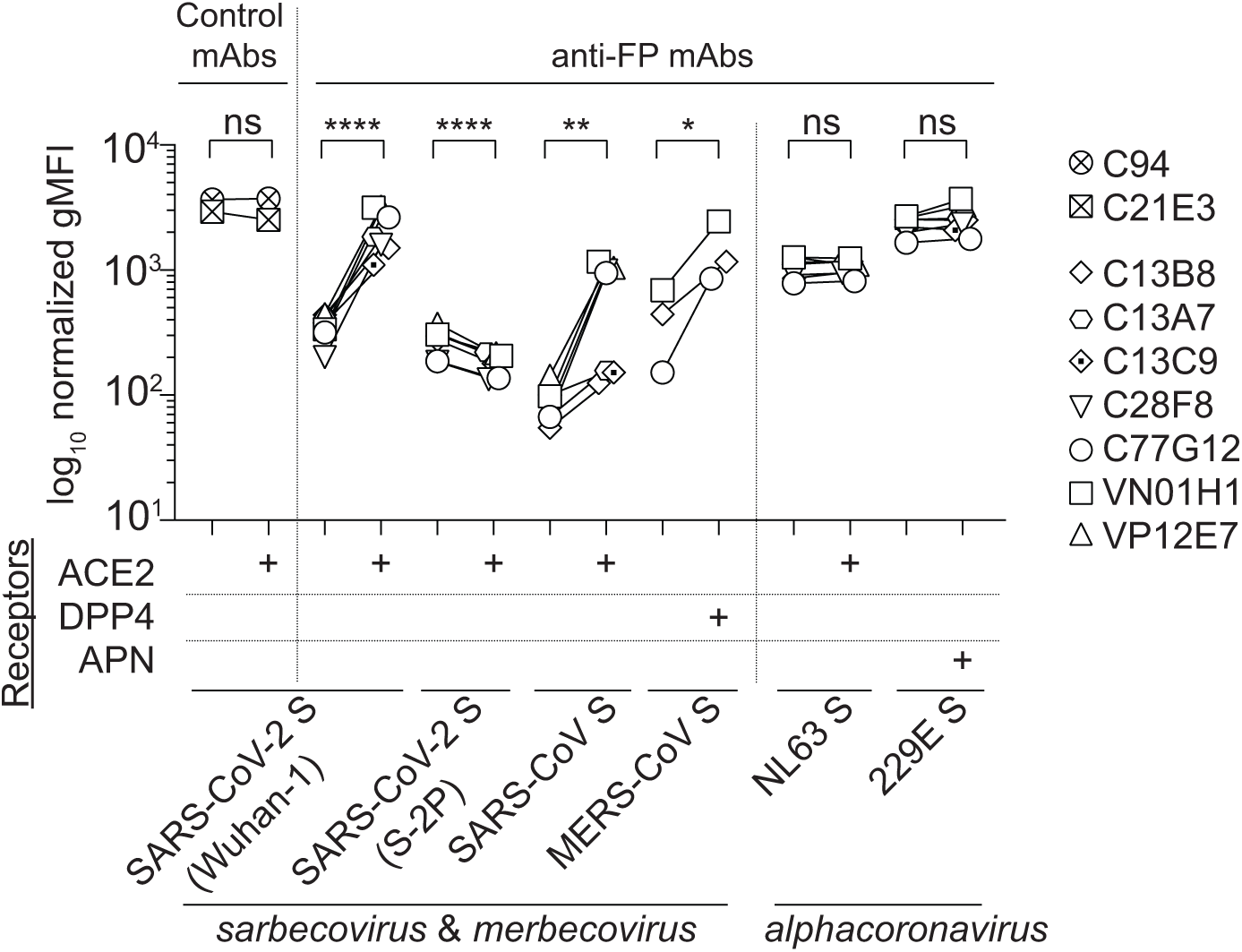
Receptor binding to SARS-CoV-2, SARS-CoV and MERS-CoV S proteins induces a conformational change that unmasks the fusion peptide epitope. Binding of anti-fusion peptide mAbs (8 μg/ml) to 293T transiently co-transfected with plasmids encoding ZsGreen and SARS-CoV-2 S (293T-S) and SARS-CoV-2 S-2P (293T-S-2P), SARS-CoV^Δ19^ S, MERS-CoV S, NL63 S and 229E S, in the presence or absence of receptors ACE2 (27 μg/ml) for SARS-CoV-2 and SARS-CoV S, DPP4 (27 μg/ml) for MERS-CoV S, and APN (27 μg/ml) for 229E S, as measured by flow cytometry. \**p* < 0.05, \*\**p* < 0.01, \*\*\*\**p* < 0.0001 Ratio paired t test.

**Table S1.**
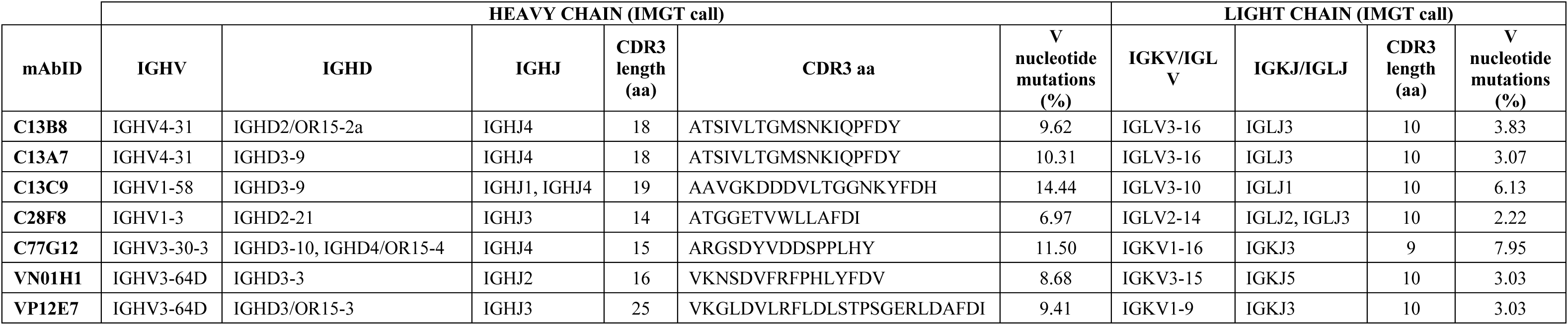
Specificity and V gene usage of the seven pan-coronavirus mAbs isolated.

**Table S2.** X-ray crystallography data collection and refinement statistics.

|  | VN01H1+peptide | VP12E7+peptide | C13C9+peptide | C13B8+peptide | C77G12+peptide |
| --- | --- | --- | --- | --- | --- |
| Space group | C 121 | P 1 | P 1 | P 212121 | P 21 |
| *Cell dimensions* |  |  |  |  |  |
| a, b, c (Å) | 152.92, 60.71, 61.11 | 71.15, 81.36, 95.65 | 63.9, 65.7, 71 | 43.3 69.3 150.7 | 45.23, 64.06, 78.8 |
| $\alpha, \beta, \gamma$ (°) | 90, 93.62, 90 | 90.34, 95.96, 100.65 | 96.5, 108.06, 110.52 | 90 90 90 | 90, 93.3, 90 |
| Resolution range(Å) | 46.24-1.86 (1.90-1.86) | 48.2-2.5 (2.56-2.50) | 44.09-2.1 (2.16-2.1) | 41.59-2.1 (2.18-2.1) | 49.67-1.7 (1.73-1.7) |
| CC1/2 | 0.998 (0.367) | 0.992 (0.737) | 0.995 (0.661) | 0.994 (0.513) | 0.999 (0.783) |
| Rmerge | 0.077 (1.6) | 0.1 (0.662) | 0.061 (0.310) | 0.131 (1.117) | 0.039 (0.497) |
| I/$\sigma$(I) | 6.5 (0.5) | 5.0 (1.3) | 8.7 (2.3) | 7.7 (1.2) | 16.4 (1.9) |
| Completeness (%) | 99.7 (99.7) | 97.9 (97.6) | 97.5 (96.6) | 93.6 (95.9) | 99.8 (99.7) |
| Redundancy | 3.3 (3.3) | 1.8 (1.9) | 2.0 (2.0) | 4.0 (4.0) | 3.7 (3.4) |
| **Refinement** |  |  |  |  |  |
| No. reflections | 46944 | 70930 | 56695 | 26948 | 49412 |
| Rwork/Rfree | 20.7/23.4 | 21.2/25.2 | 17.5/20.3 | 20.4/23.9 | 16.3/18.6 |
| *N° of atoms* |  |  |  |  |  |
| Protein | 3389 | 13636 | 6789 | 3561 | 3463 |
| Water | 285 | 535 | 808 | 244 | 517 |
| B factor (Wilson plot) | 30.91 | 38.45 | 26.53 | 30.87 | 21.08 |
| *R.m.s. deviations* |  |  |  |  |  |
| Bond lengths (Å) | 0.006 | 0.003 | 0.004 | 0.003 | 0.005 |
| Bond angles (°) | 0.85 | 0.6 | 0.72 | 0.65 | 0.82 |
| Ramachandran favored (%) | 98.43 | 97.7 | 97.96 | 98.41 | 99.32 |
| Ramachandran allowed (%) | 1.57 | 2.3 | 2.04 | 1.59 | 0.68 |
| Ramachandran outliers (%) | 0 | 0 | 0 | 0 | 0 |

- Data in parentheses are for the highest resolution shell
- $R_{\text{merge}} = \sum (\sum |I_i - \langle I \rangle| / \sum |I|)$ , where the first $\sum$ is the sum over all reflections, and the second $\sum$ is the sum over all measurements of a given reflection, with $I_i$ being the $i$ th measurement of the intensity of the reflection and $\langle I \rangle$ the average intensity of that reflection.
- $R_{\text{work}}/R_{\text{free}} = \sum (|F_o| - \langle |F_c| \rangle) / \sum |F_o|$ , where $\langle |F_c| \rangle$ is the expectation of $|F_c|$ under the probability model used to define the likelihood function. The sum is overall reflections.

**Table S3.**
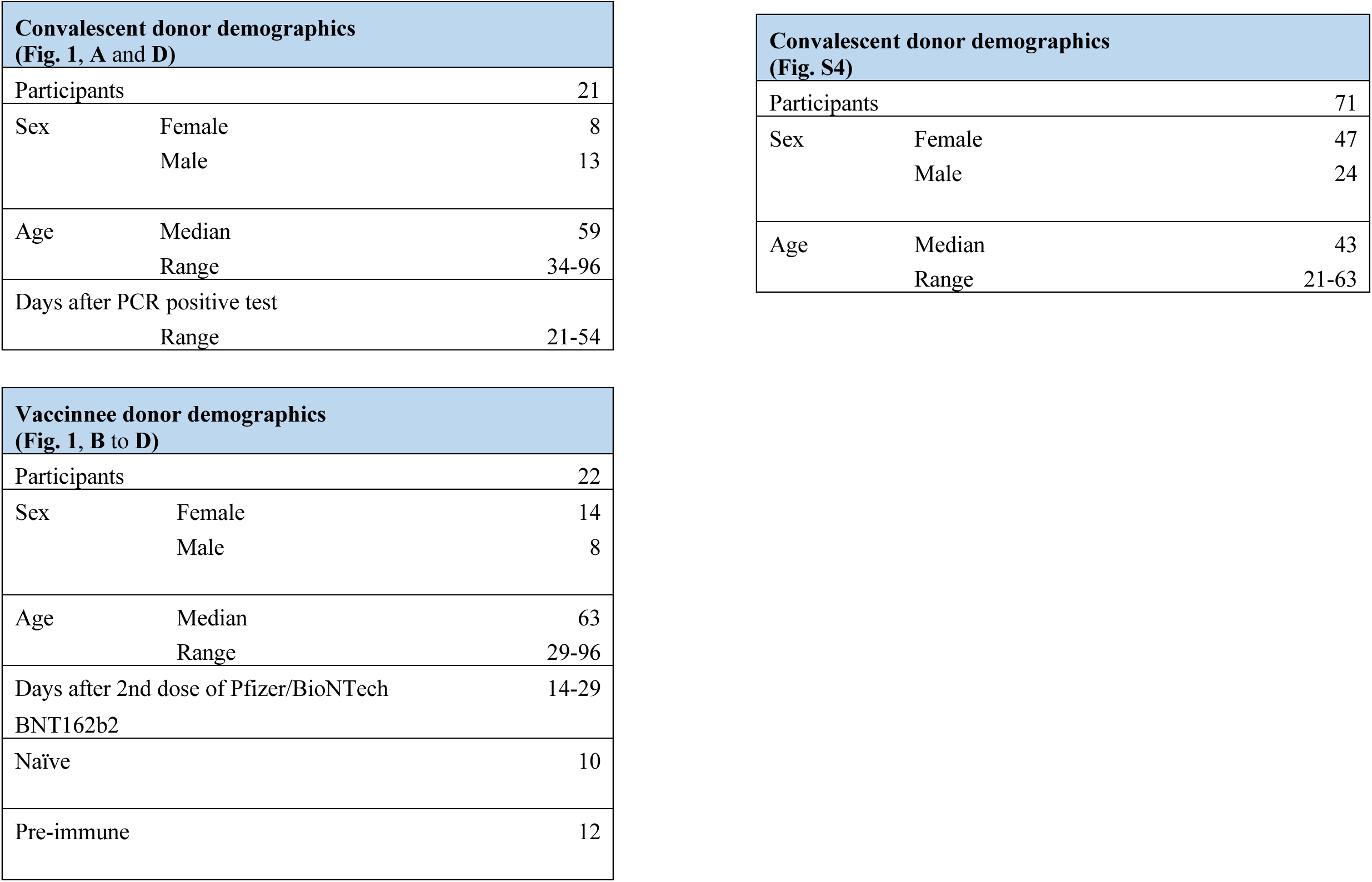
Demographics of study participants.

**Table S4.**
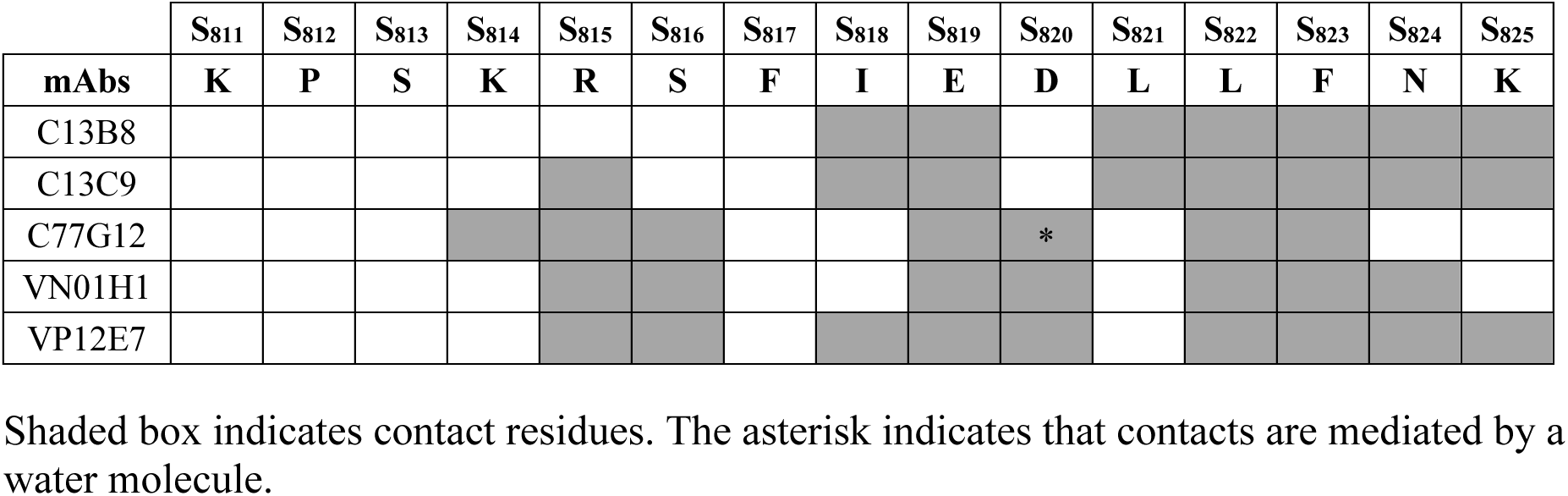
Epitope residues for each mAb as defined based on the crystal structures described in this study.

## Materials and Methods

### Samples and donors

Convalescent samples were obtained from individuals with prior SARS-CoV-2 infection, validated either by positive PCR test or by anti-spike IgG serology. Samples from vaccinated individuals (naïve or pre-immune) were collected 14-29 days after the second booster shot of Pfizer/BioNTech BNT162b2 vaccine (**Table S3**). The study protocols were approved by the Cantonal Ethics Committee of Ticino, Switzerland (CE-TI-3428, 2018-02166; CE-TI-3687, 2020-01572). All blood donors provided written informed consent for participation in the study. Human primary cell protocols were approved by the Federal Office of Public Health (no. A000197/2 to F.S.).

### Protein expression and purification

For monoclonal antibodies and Fab expression and purification, Expi293 (Gibco) cells were transiently transfected with heavy and light chain expression vectors, as previously described (*63*). The mAbs S309 and S2X58 were produced by Vir Switzerland using a similar protocol (*5, 6*). Affinity purification was performed on ÄKTA Pure 25 (Cytiva) operated by UNICORN 6.4, using HiTrap Protein A columns (Cytiva, catalog no. GE17-5079-01) for human IgG1 and CaptureSelect™ CH1-XL (Thermo Fisher, catalog no. 494346201) for Fab fragments. Buffer exchange to PBS was performed with a HiPrep 26/10 Desalting (Cytiva, catalog no. GE17-5087-01). The final products were sterilized by filtration through 0.22 μm filters and stored at 4°C. SARS-CoV-2 S HexaPro was expressed and purified as previously described (*9*). Briefly, 200 ml of Expi293F cells at 3 × 10^6^/ml were transiently transfected with 600 µl of PEI and 200 µg of the respective plasmids, following the manufacturer’s instructions. Four days post-transfection, supernatants were clarified by centrifugation at 800 g for 10 min, supplemented with 20 mM imidazole, 300 mM NaCl and 25 mM Tris-HCl pH 8.0, further centrifuged at 14,000 g for 30 min and passed through a 1 ml His trap HP column (Cytiva) previously equilibrated with binding buffer (25 mM Tris pH 7.4 and 350 mM NaCl). Proteins were eluted using a linear gradient of 500 mM imidazole using an AKTA Xpress FPLC (Cytiva) operated by UNICORN software version 5.11 (Build 407). SARS-CoV-2 S PentaPro gene, codon optimized for mammalian expression, was synthesized by GeneScript and cloned into pcDNA3.1 (-) expression vector. SARS-CoV-2 S PentaPro was expressed and purified following the same protocol as described for SARS-CoV-2 S HexaPro. SARS-CoV-2 S_2_ in postfusion conformation was produced as previously described (*11*). Briefly, SARS-CoV-2 D614G S ectodomain was produced in Expi293F cells grown in suspension using Expi293 expression medium (Life Technologies) at 37°C in a humidified 8% CO_2_ incubator rotating at 130 rpm. Cell cultures at 3 × 10^6^/ml were transiently transfected using Expifectamine (Life Technologies) following manufacturer’s instructions. Three days post-transfection, supernatant was clarified and the S glycoproetin ectodomain was affinity purified using a 1 ml HisTrapHP column (Cytiva). To isolate postfusion, S glycoprotein ectodomain was incubated with the S2X58 triggering Fab (*6*) and 1 μg/ml trypsin. After 1 h incubation, the reaction mixture was further purified using Superose 6 Increase 10/24 column (Cytiva). Purified SARS-CoV-2 S_2_ in postfusion conformation, was concentrated, buffer exchanged into PBS (137 mM NaCl, 2.7 mM KCl, 10 mM Na_2_HPO_4_, and 1.8 mM KH_2_PO_4_) and quantified using absorption at 280 nm. The gene to express SARS-CoV-2 S_2_ PentaPro in prefusion conformation was synthetized by GenScript residues 686 to 1211 fused C-terminally to a foldon trimerization domain and His-tagged. SARS-CoV-2 S_2_ PentaPro protein was expressed and purified from 200 ml of Expi293F cells. Expi293F cells grown to a density of 3 × 10^6^/ml were transiently transfected with 640 µl of Expifectamine (Thermo Fisher) and 200 µg of the corresponding expression plasmid. Cells were incubated at 37°C in a humidified 5% CO_2_ incubator rotating at 130 rpm. Four days post-transfection, supernatants were clarified by centrifugation at 800 g for 10 min, supplemented with 350 mM NaCl and 25 mM Tris-HCl pH 8.0, further centrifuged at 14,000 g for 30 min and passed through a 1 ml His trap HP column (Cytiva) previously equilibrated with binding buffer (25 mM Tris pH 7.4 and 350 mM NaCl). SARS-CoV-2 S_2_ PentaPro was eluted using a linear gradient of 500 mM imidazole. Purified protein was concentrated, buffer exchanged into TBS (25 mM Tris pH 8, 150 mM NaCl), and quantified using absorption at 280 nm.

### Enzyme-linked immunosorbent assay (ELISA)

Costar 96-well half-area plates with high protein binding treatment (Corning, catalog no. 3690) or custom made 384-well high-binding plates (Perkin Elmer) were coated overnight at 4°C with 2.5 μg/ml of the following recombinant proteins (in PBS) purchased from Sino Biological Inc: SARS-CoV-2 (2019-nCoV) S protein (S_1_+S_2_ ECD, catalog no. 40589-V08B1), SARS-CoV (S577A, Isolate Tor2) S protein (S_1_+S_2_ ECD, catalog no. 40634-V08B), MERS-CoV S protein (S_1_+S_2_ ECD, catalog no. 40069-V08B), HCoV-OC43 S protein (S_1_+S_2_ ECD, catalog no. 40607-V08B), HCoV-HKU1 (isolate N5) S protein (S_1_+S_2_ ECD, catalog no. 40606-V08B), HCoV-NL63 S protein (S_1_+S_2_ ECD, catalog no. 40604-V08B), HCoV-229E S protein (S_1_+S_2_ ECD, catalog no. 40605-V08B), SARS-CoV-2 (2019-nCoV) S_1_-His protein (catalog no. 40591-V08H), SARS-CoV-2 (2019-nCoV) S_2_ ECD-His protein (catalog no. 40590-V08B), SARS-CoV-2 (2019-nCoV) RBD-His (catalog no. 40592-V08H), Influenza A H1N1 (A/California/4/2009) Hemagglutin HA-His (catalog no. 11055-V08B1); or with 8 µg/ml of synthetic peptides of indicated sequences: TPPIKDFGGFNFSQI, DFGGFNFSQILPDPS, NFSQILPDPSKPSKR, LPDPSKPSKRSFIED, KPSKRSFIEDLLFNK, SFIEDLLFNKVTLAD, LLFNKVTLADAGFIK, VTLADAGFIKQYGDC, AGFIKQYGDCLGDIA for epitope mapping; or with a peptide mixtures of the indicted sequences: KPSKRSFIEDLLFNK (SARS-CoV-2), KPTKRSFIEDLLFNK (SARS-CoV), SRSARSAIEDLLFDK (MERS-CoV), KASSRSAIEDLLFDK (OC43), GSSSRSLLEDLLFNK (HKU1), RIAGRSALEDLLFSK (NL63), RVAGRSAIEDILFSK (229E); or with 1 µg/ml of SARS-CoV-2 prefusion and postfusion proteins. Plates were subsequently washed and blocked with Blocker Casein in PBS (Thermo Fisher Scientific, catalog no. 37528) supplemented with 0.05% Tween 20 (Sigma Aldrich, catalog no. 93773) for 1 h at room temperature. For primary antibody incubations, in serological ELISA, the coated plates were incubated with 25 µl of serial 1:3 dilutions of human plasma (12-point dilutions starting at 1:20 in Casein), for 1 h at room temperature; in cellular profiling ELISA, the coated plates were incubated with 5 µl of undiluted supernatant for 1 h at room temperature. The plates were then washed with PBS containing 0.1% Tween-20 (PBS-T), and Alkaline Phosphatase-conjugated Goat Anti-Human IgG (dilution 1:500, catalog no. 2040-04) from Southern Biotech was added and incubated for 45 min at room temperature. Plates were washed three times with PBS-T, and 4-NitroPhenyl Phosphate (pNPP, Sigma-Aldrich, catalog no. N2765) substrate was added and the absorbance of 405 nm was measured by a microplate reader (BioTek). ED50 (serum dilution) values were extrapolated by nonlinear regression curve fit (4PL) using GraphPad Prism 9 software. For PentaPro and HexaPro ELISA, 30 μl of SARS-CoV-2 S-5P or 6P at 5 ng/µl were plated onto 384-well Nunc Maxisorp (Thermo Fisher) plates in PBS and sealed overnight at room temperature. The next day, plates were washed 4 × in Tris Buffered Saline Tween (TBST) using a plate washer (BioTek) and blocked with Casein for 1 h at 37°C. Plates were washed 4 × in TBST and 1:4 serial dilutions of the corresponding mAbs starting from 0.3 mg/ml were made in 30 μl TBST, added to the plate and incubated at 37°C for 1 h. Plates were washed 4 × in TBST and 30 μl of anti-human (Invitrogen) horseradish peroxidase-conjugated antibodies diluted 1:5,000 was added to each well and incubated at 37°C. After 1 h, plates were washed 4 × in TBST and 30 μl of TMB (SeraCare) was added to every well for 5 min at room temperature. For SARS-CoV-2 S_2_ PentaPro in prefusion and SARS-CoV-2 S_2_ in postfusion conformation, the same protocol was followed except that 30 µl at 3 ng/µl of purified proteins were plated onto 384-well Nunc Maxisorp plates in TBS and sealed overnight at 4°C. Reactions were quenched with the addition of 30 μl of 1 N HCl. Plates were immediately read at 450 nm on a BioTek plate reader and data plotted and fit in Prism 9 (GraphPad) using nonlinear regression sigmoidal, 4PL, X is the concentration to determine EC50 values from curve fits.

### Cellular profiling of memory B cell repertoire analysis

Peripheral blood mononuclear cells (PBMCs) were isolated by Ficoll-Paque Plus (Cytiva, catalog no. 17-1440-03) gradient and seeded in replicative cultures of 96-well U-bottom plates (Corning, catalog no. 3799) at 10^4^ cells/well or 3 × 10^4^ cells/well. Cells were stimulated with 2.5 µg/ml of R848 (Invivogen, catalog no. tlrl-r848-5) in complete medium (RPMI 1640 medium (catalog no. 31870-025) supplemented with 2 mM glutamine (catalog no. 35050-038), 1% (vol/vol) nonessential amino acids (catalog no. 11140050, 1% (vol/vol) sodium pyruvate (catalog no. 11360-039), PenStrep (50 U/ml penicillin, 50 μg/ml streptomycin, catalog no. 15070-063), Kanamycin (50 U/ml, catalog no. 15160-047), 0.1% Beta-Mercaptethanol (catalog no. 31350-010) (all from Gibco), 10% Hyclone (Cytiva, catalog no. SH30070.03), 0.5% Transferrin (30 μg/ml) (LubioScience, catalog no. 0905-100), and 1,000 U/ml IL-2 (produced in house from transfected J558L cells). Cells were cultured at 37°C, 5% CO_2_ for six days and 5 µl of undiluted supernatant were used for ELISA (primary screening). Cultures containing antibodies with binding-profile-of-interest were selected, washed with MACS buffer (PBS with 5% FBS and 2 mM EDTA), and stained with CD19-PE-Cy7 (BD, catalog no. 341113, 1:100), IgM-AF647 (Jackson Immuno, catalog no. 109-606-129, 1:500) and IgA-AF488 (Jackson Immuno, catalog no. 109-546-011, 1:500). IgG^+^ memory B cells were isolated by negative gating strategy as CD19^+^ IgM^−^ IgA^−^ and cloned by limiting dilutions at 0.7 cells/well in complete medium. Two days later, the supernatants of the clones were screened by secondary screening ELISA to validate the binding profile.

### BCR retrieval and molecular cloning

Clones with binding-profile-of interest after secondary screening were selected and their VH and VL sequences were obtained by reverse transcription PCR (RT-PCR) using Superscript III (Thermo Fisher, catalog no. 18080044) according to manufacturer’s instruction, using the following VH, VK and VL-specific RT primers: HuIgG-const-anti 5’-TCTTGTCCACCTTGGTGTTGCT-3’; Hu-CK 5’-ACACTCTCCCCTGTTGAAGCTCTT-3’ and Hu-CL 5’-ACTGTCTTCTCCACGGTGCT-3’. VH and VK/L sequences were amplified in two nested PCR reactions using adapted VH/VK/VL primers (*64*). To facilitate high-throughput molecular cloning of BCR sequences into the VH/VK/VL expression vectors, the internal primers contain complementary 30 nucleotides of the respective expression vectors in order to be ligated by Gibson reaction using NEBuilder® HiFi DNA Assembly Master Mix (New England Biolabs, catalog no. E2621L). Ligated products were transformed into One Shot Stbl3 competent *E. coli* (Thermo Fisher, catalog no. C737303). Cloned VH, VK and VL plasmids were validated by Sanger sequencing.

### Inference for unmutated common ancestor (UCA)

Data were processed and analyzed using the Immcantation Framework (http://immcantation.org) with Change-O v1.0.2. First, the sequences were annotated using IgBlast version 1.16 (*65*) and IMGT as reference sequences (*66*). Second, clones were assigned based on IGHV genes, IGHJ gene, and junction distance with the Change-O DefineClones function. Germlines were then reconstructed using the Change-O CreateGermlines function. Finally, the phylogenic trees of each clone with their complete UCA sequences were generated with Igphyml (*67*).

### Negative-stain electron microscopy

The SARS-CoV-2 S (D614G) ectodomain trimer beginning at Q14 with a mutated S_1_/S_2_ cleavage site (SGAR), and finishing at residue K1211, followed by a TEV cleavage, fold-on trimerization motif, and an 8× His tag was expressed and purified and described previously (*68*). Three micromolar S were incubated with 3 µM of the corresponding fusion peptide Fab protein for 1 h at room temperature prior the addition of 2.6 µM of S2X58 Fab. Incubation was continued at room temperature for 1 h after which samples were diluted to 0.01 mg/ml immediately before protein was adsorbed to glow-discharged manually carbon-coated copper grids for ∼30 s before 2% uranyl formate staining. Micrographs were recorded using the Leginon software (*69*) on a 100kV FEI Tecnai G2 Spirit with a Gatan Ultrascan 4000 4k × 4k CCD camera at 67,000 nominal magnification. The defocus ranged from 2.0 to 4.0 μm and the pixel size was 1.6 Å.

### Epitope substitution scan

Epitope substitution scan was performed by PEPperPRINT GmbH, Heidelberg, Germany. Briefly, each amino acid in the fusion peptide sequence K_811_PSKRSFIEDLLFNKVTLAD_830_ was substituted stepwise with all 20 main amino acids. These peptide variants and wildtype peptide, as well as HA control peptides was printed on a microarray chip in triplicate. Primary antibodies at 1 µg/ml (C13B8 and C13C9) or 100 µg/ml (VN01H1 and VP12E7) were incubated with the microarray chip for 16 h at 4°C with orbital shaking at 140 rpm. After washing, secondary antibody goat anti-human IgG (H+L) DyLight680 (0.2 µg/ml) was incubated for 45 min at room temperature before reading on Innopsys InnoScan 710-IR Microarray Scanner.

### Generation of recombinant ACE2-mFc

Residues 18-615 of human ACE2 (UniProtKB - Q9BYF1) were synthesized by Genscript and cloned into pINFUSE-mIgG2b-Fc2 expression plasmid (InvivoGen). Recombinant protein was produced by transient transfection of Expi293 cells and purified using HiTrap Protein A column. Buffer exchange was performed using HiPrep 26/10 Desalting column and final product was sterilized through a 0.22 μm filter.

### Cell lines and media culture

293T and A549 cells were cultured in high glucose DMEM (Gibco, catalog no. 61965-026) supplemented with 10% FBS, 1% (vol/vol), nonessential amino acids, 1% (vol/vol), sodium pyruvate and PenStrep (50 U/ml penicillin, 50 μg/ml streptomycin). HuH-7 cell line was obtained from JCRB Cell Bank and cultured in DMEM (Gibco, catalog no. 31885-023) supplemented with 10% FBS, 1% (vol/vol), nonessential amino acids, 2 mM glutamine and PenStrep (50 U/ml penicillin, 50 μg/ml streptomycin). Vero-TMPRSS2 cells were cultured in DMEM (Gibco, catalog no. 11995-040) supplemented with 10% FBS (VWR, catalog no. 97068-085) and PenStrep (Gibco, catalog no. 15140-122) (*35*).

### Generation of overexpression cell lines

pLVX-puro-ACE2 transfer plasmid was kindly provided by Manfred Kopf (ETH Zurich). pLVX-EF1a-TMPRSS2-IRES-ZsGreen1 transfer plasmid was generated from the reference pWPI-IRES-Bla-Ak-TMPRSS2 plasmid (Addgene, catalog no. 154982). pLVX-puro-spike transfer plasmid was generated from reference pHDM-SARS-CoV-2 spike (BEI resources, catalog no. NR-52514). Stable cell lines were generated using VSV-based lentivirus transduction. Briefly, 293T cells at 70-80% confluency in T75 flask were co-transfected with transfer plasmids encoding genes of interest (ACE2, TMPRSS2 or SARS-CoV-2 S) and packaging plasmid psPax2 and envelope plasmid pMD.2G with polyethylenimine (PEI) (at a ratio of 1:2.3 DNA:PEI) (Polysciences, catalog no. 24765-2) in Optimem (Life Technologies Europe BV, catalog no. 31985047). Supernatants containing lentiviral particles were harvested 36 h post-transfection, filtered through 0.22 μm filter and precipitated using 40% (W/V) PEG-8000 (Promega, catalog no. V3011) and 1.2M NaCl (Sigma-Aldrich, catalog no. 71380) for 4-6 h on a shaker at 4°C, and thencentrifuged for 1 h at 1,600 g at 4°C. The lentivirus-containing pellet was resuspended in 100 μl media and was used to transduce 293T or A549 cell lines. 293T-ACE2, 293T-S, A549-ACE2 and A549-S cell lines were selected using 10 μg/ml puromycin (InvivoGen, catalog no. ant-pr-1) 4 days post-transduction. 293T-ACE2-TMPRSS2-GFP cell line was generated from 293T-ACE2 cells by subsequent transduction of pLVX-EF1a-TMPRSS2-IRES-ZsGreen1-containing lentivral prep and sorted using BD FACSAria III. A549-ACE2-TMPRSS2 and Huh7-TMPRSS2 stable cell lines were generated using commercial lentivirus (Addgene, catalog no. 154982-LV) and selected using 10 μg/ml blasticidin (InvivoGen, catalog no. ant-bl-1) 4 days post-transduction.

### Pseudotyped virus production

To produce SARS-CoV-2, SARS-CoV, MERS-CoV and 229 S pseudotyped virus, full-length spike-encoding plasmids were obtained from the following manufacturers; SARS-CoV-2 Wuhan-Hu-1 (catalog no. NR-52514) from Bei Resources; SARS-CoV-2 Omicron BA.1 (VG40835-UT), SARS-CoV (VG40150-G-N), MERS-CoV (VG40069-G-N) and 229E (VG40605-UT) from SinoBiological; NL63 (YP_003767.1) and WIV-1 (Uniprot – U5WI05) were synthesized from Genscript. HIV-based HcoV spike glycoprotein-pseudotyped viruses were prepared as previously described (*70*) with slight modifications. Briefly, 293T cells were co-transfected with a lentiviral backbone encoding luciferase reporter (pHAGE-CMV-Luc2-IRES-ZsGreen-W (Bei Resources, catalog no. NR-52516), HIV-based packaging plasmids (Tat, Gag-Pol and Rev) (Bei Resources, catalog no. NR-52518, NR-52517 and NR-52519) and various spike expression plasmids using PEI in Optimem. Supernatants were harvested 36 h post-transfection and pseudotyped viral particles were precipitated as described above. To produce NL63 VSV virus, HEK-293 cells were transfected with a pcDNA3.1 expression vector encoding full-length S harboring a truncation of the 20 C-terminal residues to improve membrane transport. The day after transfection, cells were transduced with VSVΔG/Luc (*71*). After 2 h, infected cells were washed four times with DMEM before adding medium supplemented with anti-VSV-G antibody (I1- mouse hybridoma supernatant diluted 1 to 25, from CRL-2700, ATCC). Supernatant was harvested 18-24 h post inoculation, clarified from cellular debris by centrifugation at 2,000 g for 5 min, filtered using a 0.45 µm membrane, concentrated 10 times using a 30 kDa cut off membrane (Amicon), aliquoted and frozen at -80°C until use.

### Pseudotyped virus neutralization

For SARS-CoV-2, SARS-CoV, MERS-CoV, 229E and WIV-1 S pseudotyped virus neutralization assay, target cells (293T-ACE2 or 293T-ACE2-TMPRSS2 for SARS-CoV-2 and SARS-CoV; HuH-7-TMPRSS2 for MERS-CoV and 229E; 293T-ACE2-TMPRSS2 for WIV-1) were seeded in white 96-well plate (Perkin Elmer, catalog no. 6005688) at 40,000 cells/well the day before infection. Concentrated viruses were titrated in serial dilutions with the respective target cell lines and the luciferase reporter signal was determined 48 h later using Luciferase Assay System (Promega, catalog no. E1501) on Cytation 3 (BioTek). Virus concentrations that gave signal higher than 10^5^ RLUs/well were used in neutralization experiments. Serial 1:3 dilutions of mAbs (10- point dilutions starting at 200 μg/ml) were pre-incubated with pseudotyped viruses at 37°C for 30 min. The pseudotyped virus-mAb mixture were then overlayed onto target cell lines in the presence of 5 μg/ml polybrene (Sigma Aldrich, catalog no. TR-1003-G) and analyzed 48 h post-infection. For NL63 S pseudotyped virus neutralization assay, Vero E6-TMPRSS2 cells maintained in DMEM supplemented with 10% FBS and 1% PenStrep, were seeded into white 96-well plates at 45,000 cells/well and cultured overnight at 37°C. Eleven-point 3-fold serial dilutions of the corresponding mAbs were prepared in DMEM. NL63 S pseudotyped viruses were added 1:1 (v/v) to each dilution (final volume 50 μl) and the mixtures were incubated at 37°C. After 45-60 min incubation, 40 μl of each reaction mixture were added to the cells which were incubated at 37°C. After 2 h incubation, 40 μl DMEM were added to avoid evaporation and incubation was continued at 37^°^C. After 17-20 h, 60 μl/well of One-Glo-EX substrate (Promega) were added to the cells and incubated in the dark for 5-10 min prior reading on a Varioskan LUX plate reader (Thermo Fisher). Data was processed using GraphPad Prism v9.0.

### Inhibition of cell-to-cell fusion

For testing inhibition of spike-mediated cell–cell fusion, A549-S and A549-ACE2-TMPRSS2 cells were stained with CFSE (Thermo Fisher, catalog no. C1157) and CellTrace™ Far Red (Thermo Fisher, catalog no. C34572), respectively, according to manufacturer’s instruction. Stained cells were resuspended in complete media containing Hoechst 33342 (Thermo Fisher, catalog no. H1399) at a final concentration of 5 μg/ml. A549-S cells were co-cultured in indicated concentrations of mAbs for 30 min at 37°C before addition of stained-A549-ACE2-TMPRSS2. Fusion events were measured 2 h post incubation with Molecular Devices ImageXpress Micro 4 system. Acquisition was performed with a 20x/0.45 Super Plan Fluor ELWD objective, FITC and Cy5 filter and images collected with a Andor Zyla sCMOS camera. Nine fields per well were imaged and were subsequently processed with Metaxpress and Powecore softwares.

### SARS-CoV-2 infection model in hamsters

KU LEUVEN R&D has developed and validated a SARS-CoV-2 Syrian Golden hamster infection model (*72, 73*). This model is suitable for the evaluation of the potential antiviral activity and selectivity of novel compounds/antibodies (*74*). The SARS-CoV-2 strain used in the model, Gamma P.1 (EPI_ISL_1091366; 2021-03-08), was recovered from a nasopharyngeal swab taken from a traveler returning to Belgium in March 2021 (*75*). The variant was subjected to sequencing on a MinION platform (Oxford Nanopore) directly from the nasopharyngeal swabs; passage 2 virus on Vero E6 cells was used for the study described here. The titer of the virus stock was determined by end-point dilution on Vero-E6 cells by the Reed and Muench method (*76*). Live virus-related work was conducted in the high-containment A3 and BSL3+ facilities of the KU Leuven Rega Institute (3CAPS), under licenses AMV 30112018 SBB 219 2018 0892 and AMV 23102017 SBB 219 20170589, according to institutional guidelines.

Syrian Golden hamsters (Mesocricetus auratus) were purchased from Janvier Laboratories and were housed per two in ventilated isolator cages (IsoCage N Biocontainment System, Tecniplast) with ad libitum access to food and water and cage enrichment (wood block). The animals were acclimated for 4 days prior to study start. Housing conditions and experimental procedures were approved by the ethics committee of animal experimentation of KU Leuven (license P065-2020). Female hamsters of 6-8 weeks old were anesthetized with ketamine/xylazine/atropine and inoculated intranasally with 50 µl containing 1×10^4^ TCID50 SARS-CoV-2 gamma variant (day 0). Animals were prophylactically treated 24 h before infection with VN01H1 (50 mg/kg), C77G12 (25 mg/kg and 50 mg/kg) and MGH2 (50 mg/kg) via intraperitoneal (IP) administration. Hamsters were monitored for appearance, behavior and weight. At day 4 post infection (pi), hamsters were euthanized by IP injection of 500 μl Dolethal (200 mg/ml sodium pentobarbital, Vétoquinol SA). Lungs were collected and viral RNA and infectious virus were quantified by RT-qPCR and end-point virus titration, respectively. Blood samples were collected for pharmacokinetics analysis.

### SARS-CoV-2 RT-qPCR

Hamster lung tissues were collected after sacrifice and were homogenized using bead disruption (Precellys) in 350 µL TRK lysis buffer (E.Z.N.A.^®^ Total RNA Kit, Omega Bio-tek) and centrifuged (10.000 rpm, 5 min) to pellet cell debris. RNA was extracted according to the manufacturer’s instructions. Of 50 μl eluate, 4 μl was used as a template in RT-qPCR reactions. RT-qPCR was performed on a LightCycler96 platform (Roche) using the iTaq Universal Probes One-Step RT-qPCR kit (BioRad) with N2 primers and probes targeting the nucleocapsid (*72*). Standards of SARS-CoV-2 cDNA (IDT) were used to express viral genome copies per mg tissue or per ml serum.

### End-point virus titrations

Lung tissues were homogenized using bead disruption (Precellys) in 350 µl minimal essential medium and centrifuged (10,000 rpm, 5min, 4°C) to pellet the cell debris. To quantify infectious SARS-CoV-2 particles, endpoint titrations were performed on confluent Vero E6 cells in 96- well plates. Viral titers were calculated by the Reed and Muench method (*77*) using the Lindenbach calculator and were expressed as 50% tissue culture infectious dose (TCID50) per mg tissue.

### Histology

For histological examination, the lungs were fixed overnight in 4% formaldehyde and embedded in paraffin. Tissue sections (5 μm) were analyzed after staining with hematoxylin and eosin and scored blindly for lung damage by an expert pathologist. The scored parameters, to which a cumulative score of 1 to 3 was attributed, were the following: congestion, intra-alveolar hemorrhagic, apoptotic bodies in bronchus wall, necrotizing bronchiolitis, perivascular edema, bronchopneumonia, perivascular inflammation, peribronchial inflammation and vasculitis.

### Crystallization, data collection, structure determination and analysis

Fabs C13B8, C13C9, C77G12, VN01H1 and VP12E7, at 20 mg/ml were mixed with the fusion peptide (KPSKRSFIEDLLFNK, GenScript) at 20 mM and incubated for 2 h at room temperature before setting up crystallization plates. Crystals of Fabs in complex with the fusion peptide were obtained at 22°C by sitting drop vapor diffusion. A total of 100 nl complex were mixed with 100 nl mother liquor solution containing 0.2M potassium acetate, 20% (w/v) PEG 3350, 0.2 M potassium chloride, 0.05 M HEPES-NaOH pH 7.5, 35% (v/v) pentaerythritol propoxylate 5/4/PO/OH (VN01H1 complex); 0.2 M Calcium Acetate Hydrate, 0.1 M MES NaOH, pH 6.0, 25% PEG 8000 (VP12E7 complex); 0.2 M Calcium Chloride 20 % (w/v) PEG 3350 (C77G12 complex); 0.17 M Ammonium Acetate, 0.085 M Sodium Acetate: HCl, pH 4.6, 25.5 % (w/v) PEG 4000, 15 % (v/v) glycerol (C13B8 complex) and 0.04 M KH2PO4, 16 % (w/v) PEG 8000, 20 % (v/v) Glycerol (C13C9 complex). Drops were equilibrated against reservoir solutions for 1 week at room temperature after which crystals were flash cooled in liquid nitrogen using the mother liquor solution supplemented with 30% glycerol as a cryoprotectant. Data were remotely recorded at the Molecular Biology Consortium beamline 4.2.2 at the Advanced Light Source synchrotron facility in Berkeley, CA. Individual datasets for each complex were processed with the XDS software package (*78*) and Mosfilm (*79*) and scaled using and SCALA or aimless (*80*). Initial phases were obtained by molecular replacement in Phaser (*81*) on the CCP4 suite, using crystal structures of Fabs as search models. Several subsequent rounds of model building and refinement were performed using Coot (*82*), Refmac5 (*83*) and Buster (*84*) to arrive to the final model for each complex.

### Transient expression and monoclonal antibody staining of HCoV S-expressing 293 cells

For transient expression of HCoV spike proteins, 293T cells were co-transfected, with plasmid encoding ZsGreen (Bei Resources, catalog no. NR-52516) and corresponding HCoV spike proteins: SARS-CoV-2 Wuhan-Hu-1 S (catalog no. NR-52514) from Bei Resources; MERS-CoV S (VG40069-G-N), 229E S (VG40605-UT), NL63 S (VG40604-UT) from SinoBiological; SARS-CoV S (VG40150-G-N) from SinoBiological that was cloned into pHDM expression plasmid with 19 amino-acid C-terminal truncation (*85*), using PEI in Optimem as above. For the SARS-CoV-2 S2P mutant, K986P and V987P mutations were introduced into the wildtype backbone using Q5 Site-Directed Mutagenesis Kit (NEB, catalog no. E0554S). Transiently transfected cells were stained the following day with mAbs conjugated using DyLight® 650 Conjugation Kit (Fast) - Lightning-Link (Abcam, catalog no. ab201803) according to manufacturer’s instructions. Dylight 650-conjugated mAbs (of indicated concentrations) were incubated with 50,000 un-trypsinized 293T-S cells per well in the presence or absence of ACE2-mFc (27 μg/ml), S2E12 (20 μg/ml), S2M11 (*42*) (20 μg/ml), DPP4-hFc (27 μg/ml) and APN-hFc (27 μg/ml) for 2 h at room temperature in MACS buffer. Cells were then washed and analyzed by flow cytometry using BD Symphony and FlowJo.

